# Integrated analysis of stemness-associated immune modulatory circuits in squamous cell carcinomas

**DOI:** 10.64898/2026.04.02.716182

**Authors:** Weijie Guo, Daniel Leon, Jingyun Luan, Audrey Gang, Xuejie Huang, Benjamin Nicholson, Karolina Dorosz, Yinhan (Stacey) Zhao, Sarra Grimshaw, Diana Bolotin, Mark W. Lingen, Everett E. Vokes, Alexander T. Pearson, Ari J. Rosenberg, Le Shen, Evgeny Izumchenko, Nishant Agrawal, Jean X. Jiang, Yuxuan Miao

## Abstract

Emerging evidence indicates that a subset of cancer cells enriched for stemness-related gene signatures possess distinct immunomodulatory capacities, enabling these tumor-initiating stem cells (tSCs) to more effectively evade or resist anti-tumor immunity. Despite these advances, the tSC-specific molecular circuits orchestrating their specialized immune privilege program are not well defined. Here, in squamous cell carcinomas of the skin and oral cavity, we comprehensively delineate the unique immune-evasive properties of tSCs and dissect the transcriptional regulation shaping their immunomodulatory programs. By integrating transcriptome profiling, chromatin landscape mapping, genetic perturbation, and single-cell RNA sequencing, we found that the tSC-specific immune program is broadly governed by SOX2, a stemness-associated transcription factor. We demonstrate that SOX2 enables tSCs to sustain immature tumor-associated neutrophils (TANs) and subsequently trigger these myeloid cells to foster the development of tumor-associated macrophages (TAMs). This SOX2-directed tSC-TAN-TAM axis establishes a localized immunosuppressive niche for protecting tSC.

**SIGNIFICANCE:** Here, we uncover SOX2 as a master regulator that orchestrates conserved immune modulatory circuits in tSCs to sustain pro-tumor myeloid cell states. These findings place tSCs at the apex of immune landscape remodeling, asserting a central role of stemness-associated program in organizing the immunosuppressive tumor microenvironment.

## INTRODUCTION

Escaping immune surveillance is a prerequisite for oncogenic transformation and malignant progression. Therefore, therapeutic strategies that disrupt mechanisms employed by cancer cells to evade anti-tumor immunity have revolutionized cancer treatments (1,2). However, a major limitation of current regimen is patient relapse, with tumors frequently reemerging even when initial responses appear robust (3-5). This challenge is particularly urgent for squamous cell carcinomas (SCC), which typically arise in the skin (cutaneous SCCs or CSCCs), oral cavity (head and neck SCCs or HNSCCs), lung, and cervix (6-8). Even though the development of immune checkpoint blockade (ICBs) has extended overall survival in SCC patients, especially those with recurrent/metastatic HNSCCs and CSCCs (9,10), response durability remains limited compared to other cancers (4,9). Notably, more than 60% of SCC tumors rapidly relapse after an initial response to immunotherapy (4,9).

A major reason current immunotherapies often fail is because most treatments are designed under the assumption that all tumor cells utilize similar mechanisms to evade anti-tumor immunity. Recent studies have challenged this paradigm, revealing substantial heterogeneity among cancer cell populations. In genetically engineered mouse models of CSCCs and HNSCCs, a subset of tumor-initiating stem cells (tSCs) has been shown to be refractory to cytotoxic T lymphocyte (CTL) attacks, despite intact presentation of targeted antigens (11,12). In both CSCCs and HNSCCs, these stemness-enriched cancer cells are situated predominantly at the tumor-stroma interface (13-17). By co-opting molecular features of normal epithelial stem cells, these tSCs play key roles in driving tumor formation, propagation, metastasis, and resistance (18-20). Due to their enhanced evasion from immunotherapy, while other differentiated cells can be easily killed by CTLs, elucidating the stemness-associated immune modulatory mechanisms is essential for designing novel strategies to target these key cell population in SCCs. Moreover, many solid tumors, including glioblastoma (21) and colon cancers (22), exhibit similar hierarchical organization governed by stem-like cancer cells. Even in certain cancers where no such a discrete tSC population can be clearly defined, tumor cells often acquire stemness or progenitor-like features that drive tumorigenesis. Thus, understanding the stemness-associated immune evasive mechanisms in SCCs is likely to provide applicable insights for improving immunotherapeutic strategies across diverse malignancies.

Despite the critical roles of tSCs in driving cancer relapse after immunotherapy, their unique immune evasive mechanisms have only recently begun to be elucidated. Several immune suppressive ligands, including CD80 and CD276, have been shown to be selectively enriched in tSCs, enabling these cells to directly dampen T cell functions. Moreover, tSCs can be shielded by immune suppressive cells that establish a specialized niche within the tumor microenvironment (TME) (23,24). In CSCC, for instance, tSCs have been reported to produce high level of IL33, which modulates the differentiation trajectory of a subset of FcεRIα+ tumor associated macrophages (TAMs) that develop in close proximity of tSCs (25). Furthermore, tSCs produce arachidonic acid, which enhances the immune suppressive functions of tumor-associated neutrophils (TANs) (26), thereby protecting tSCs during immunotherapy treatments.

However, due to the lack of comprehensive molecular analyses, the key regulatory mechanisms shaping stemness-associated immune resistance in tSCs remain not defined. Importantly, these tSCs represent the major tumor population responding to immune suppressive signaling, such as TGFβ (16,17,25,27). These signaling pathways can endow tSCs with specialized properties, including activation of CD80, that enhance resistance to anti-tumor immunity (11,12). Furthermore, many defining molecular features of tSCs are regulated by stem cells-specific transcription factors (TFs). Yet, it remains unclear which TF serve as a master regulator to orchestrate the diverse immune modulatory functions of tSCs. Addressing this key question is imperative not only for understanding tSC specific immune evasive mechanisms, but also for developing therapeutic strategies to broadly dismantle the tSC-specific immune resistance and prevent cancer relapse following immunotherapy.

In this study, we sought to identify the master regulator that activates and coordinates the tSC-specific immune modulatory program. To this end, we compared the transcriptome and chromatin landscape of tSCs with other tumor populations isolated from the same neoplasms. We further mapped critical TF binding profile and utilized single cell analysis to determine the impact of genetic perturbation on the immune landscape. This integrated analysis enabled the identification of conserved molecular circuits orchestrated by a stem cell specific master regulator, which organizes and sustains the immune suppressive tumor microenvironment (TME).

## RESULTS

### Dissecting the molecular features of tSCs in autochthonous squamous cell carcinoma models

To identify the stemness-associated network that conferring superior immune-evasive properties to tSCs, we compared the transcriptomes of tSCs with those of other tumor populations in mouse SCCs originating from distinct tissues, including the skin and oral cavity. Given that most cancer immunology studies to date have relied on transplanted tumor cell lines, most tSC-specific features have likely been overlooked. To capture the accurate molecular properties of tSCs, we therefore utilized autochthonous tumor models. Specifically, spontaneous CSCCs were generated using a well-established two-step chemical mutagenesis protocol in which C57/BL6 mouse skin epithelium was exposed to mutagen 7,12-dimethylbenz[*a*]anthracene (DMBA), followed by repeated applications of the phorbol ester 12-O-tetradecanoylphorbol 13-acetate (TPA) (Fig. 1A). To induce spontaneous HNSCCs, the C57/BL6 mouse oral cavity was exposed to 4-nitroquinoline 1-oxide (4-NQO), mimicking tobacco-associated carcinogenesis (Fig. 1A). Importantly, these approaches induce autochthonous SCC tumors with mutational profiles, developmental trajectories, hierarchical organization, heterogeneous compositions, and complex ecosystems that closely recapitulate human tumors. By pooling and profiling tumors from different animals, we minimized the confounding effects of gene signatures linked to specific oncogenic mutations, enabling identification of conserved stemness-associated immune-evasive programs.

**Figure 1.**
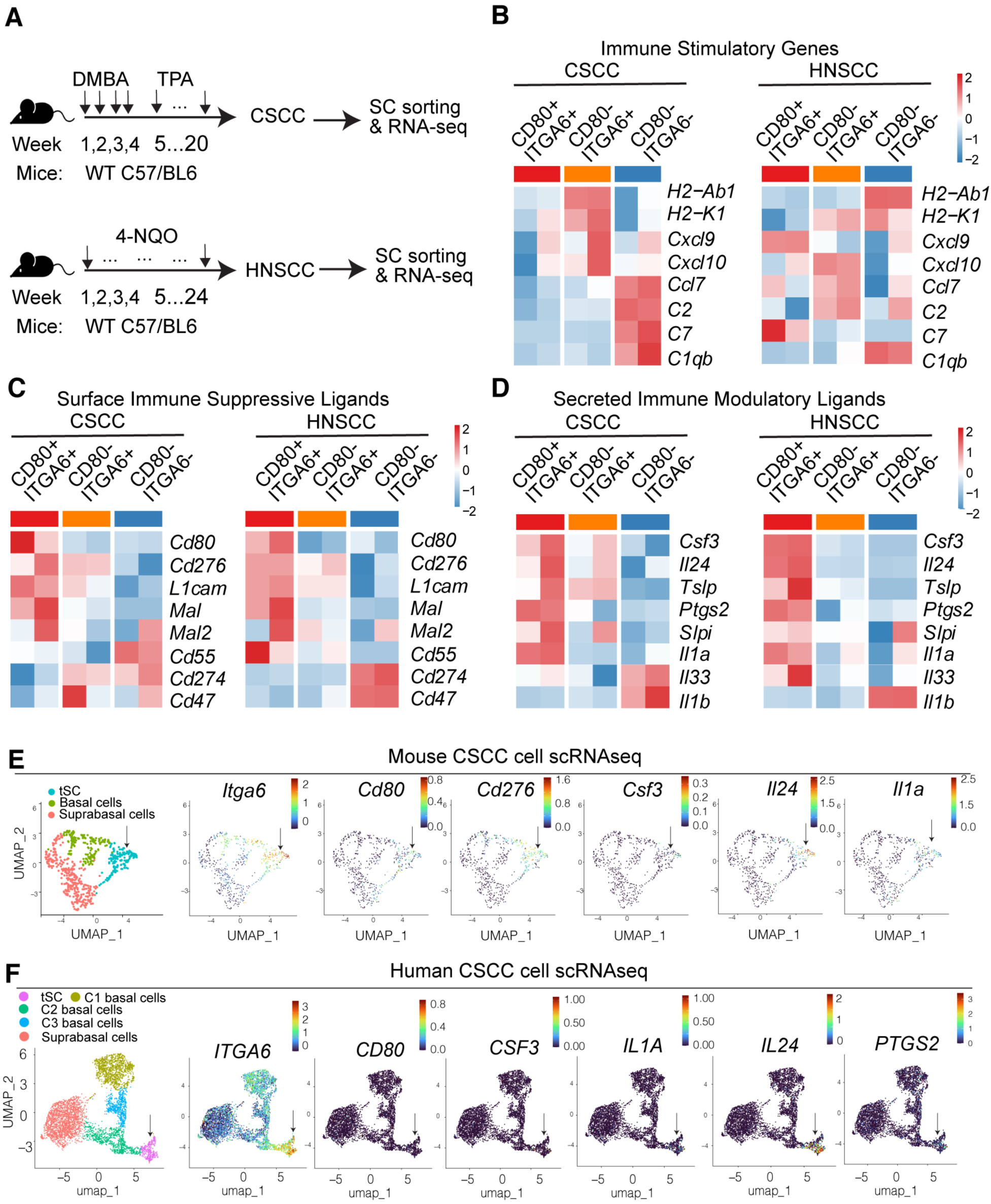
Tumor-initiating stem cells exhibit distinct immune-modulatory profile in squamous cell carcinomas. **A**, Schematic of experimental procedures for inducing spontaneous mouse CSCCs and HNSCCs for transcriptome analysis. **B-D**, Heatmap showing the differentially expressed immune stimulatory genes (**B**), immune suppressive surface ligands (**C**), and secreted immune modulatory ligands (**G**) in CD80^+^ITGA6^+^ tSCs, CD80^-^ITGA6^+^ basal cells, and CD80^-^ITGA6^-^ suprabasal cells in CSCCs and HNSCCs. **E** and **F**, UMAP showing the signature and immune modulatory genes expressed in various cancer cell clusters in mouse CSCCs (**E**) or human CSCCs (**F**).

When the tumors developed, using established surface markers, we isolated various tumor cell populations by fluorescence-activated cell sorting (FACS). Specifically, ITGA6 and CD80 were used to define and isolate TGFβ-responsive tSCs in both CSCCs and HNSCCs (Supplementary Fig. S1A and S1B). Previous limited dilution transplantation assays and lineage tracing experiments confirmed that, compared to other tumor cells from the basal layer, CD80^+^ TGFβ-responsive basal cells exhibit the highest tumor initiating potentials compared to other basal tumor cells (28,29). In parallel, we isolated CD80^-^ ITGA6^+^ basal cells and CD80^-^ ITGA6^-^ subrabasal cells from the same SCCs (Supplementary Fig. S1A and S1B), subjecting these populations to bulk RNA sequencing (RNA-seq) for transcriptomic analysis.

Differential gene expression and gene set enrichment analyses identified molecular signatures enriched in CD80^+^ITGA6^+^ tSCs compared to other tumor populations. As we expected, in SCCs from both skin and oral cavity, CD80^+^ITGA6^+^ tumor cells shared many common molecular features, including elevated expression of stem cell specific transcription factor (e.g.,*Sox2, Trp63)*, signaling molecules regulating stem cell activites, (e.g.,*Hmga1)*, as well as surface markers frequently used to identify cancer stem cells (e.g.,*Cd44)* (Supplementary Fig. S1C). Consistent with previous reports (29,30), CD80^-^ITGA6^+^ basal tumor cells also express certain stemness-related genes, although at lower levels than those observed in the CD80^+^ITGA6^+^ cells (Supplementary Fig. S1C). Together, these finding confirm the validity of our sorting strategy for separating cancer cell populations with high, intermediate, or low stemness-related features in SCCs.

### Systematic characterization of immune modulatory properties of tSCs

Having confirmed the stemness-associated signatures in sorted tSCs, we next examined gene expression patterns whose products are likely to exert profound effects on the immune microenvironment. Compared to CD80^-^ basal cells or ITGA6^-^ suprabasal cells, the double positive tSCs exhibited marked down-regulation of the antigen presentation genes such as *H2-Ab1* and *H2-K1* (Fig. 1B). In addition, tSCs in both CSCCs and HNSCCs exhibited reduced expression of pro-inflammatory chemokines (e.g., *Cxcl10*) (Fig. 1B), which are critical for recruiting anti-tumor T cells and proinflammatory myeloid cells, particularly monocytes. The selective silencing of chemokine production in tSCs (Supplementary Fig. S1D), contrasted with the elevated expression of the same chemokine cohort in other tumor cells, likely contributes to shaping the spatial distribution of various immune cells within tumors. Furthermore, genes related to complement activation were also collectively downregulated in tSCs (Fig. 1B). Notably, suppression of complement genes is more profound in CSCCs than in HNSCCs (Fig. 1B), highlighting tissue-specific differences in complement pathway function. Although both anti-tumor and pro-tumor roles of complement have been reported, the near-uniform downregulation of complement genes in tSCs from CSCCs suggests that complement-mediated T cell priming functions may be particularly important in the skin context.

Importantly, tSCs not only downregulate immune stimulatory genes but also upregulate multiple factors that can significantly influence immune cell activities. One prominent group of genes enriched in tSC encodes surface ligands (Fig. 1C). Although many surface ligands have been shown to be co-opted by cancer cells to suppress T cell responses, most of these conclusions were drawn from studies using grafted homogenous cancer cell lines. As a result, the precise expression pattern of these ligands in cancers have often been overlooked. To our surprise, we found that *Cd47 and Cd274* (gene encoding PDL1) were expressed at higher levels in differentiated cancer cells, even though these cells are preferentially targeted and eliminated by T cells (Fig. 1C) (28). Instead, tSCs selectively acquire a unique set of immune suppressive ligands, such as *Cd276 and L1cam* (Fig. 1C). While CD276 can directly suppress T cell functions (31), L1CAM has been shown to promote the expansion of regulatory T cells (32). In addition, Mal family proteins, known to inhibit antigen presentation (33), are also specifically enriched in tSCs (Fig. 1C). Together, these surface ligands allow tSCs to directly modulate immune cell behavior through cell-cell interactions.

In addition to the enriched expression of immune suppressive surface ligands, we also identified multiple secreted immune modulatory factors that enable tSCs to exert long-range effects and to orchestrate the differentiation trajectories of various immune populations. Unexpectedly, multiple cytokines that are thought to be broadly produced by all cancer cells were found to be selectively enriched in tSCs. For example, compared to CD80^-^ basal cells or suprabasal tumor cells, tSCs express the highest levels of *Tslp*, which promotes suppressive macrophage development and inhibits Th1 cell functions (Fig. 1D) (34,35). tSCs also produce *Il24* (which can stimulate Th17 cells to induce IL10 production) (36), *Slpi* (which suppresses the anti-tumor functions of neutrophils) (37), and *Ptgs2* (which directly inhibits IL2 signaling in CD8 T cells) (Fig. 1D) (38,39). The expression patterns of IL1 family cytokines are particularly notable. While *Il1β* is predominantly expressed by differentiated tumor cells, *Il1α* is strongly enriched in tSCs (Fig. 1C), suggesting a potential role of tSCs in shaping emergency myelopoiesis and supplying the TME with multiple myeloid populations (40). Consistent with this possibility, we observed marked enrichment of *Csf3* expression in tSCs, a cytokine known to orchestrate myeloid cell differentiation (Fig. 1D). Although *Csf3* has been implicated in promoting myeloid cell development, it has generally been assumed that most cancer cells are capable of producing this cytokine. In contrast, our data indicate that Csf3 production is largely restricted to tSCs, pointing to a specialized mode of communication between tSCs and myeloid cells.

### Single cell profiling of the immune modulatory gene signatures of tSCs

To further validate the expression patterns of these immune modulatory genes uncovered by bulk RNA-seq, we analyzed published singe cell RNA-seq datasets. First, we examined the single cell transcriptome of cancer cells isolated from spontaneous mouse skin SCCs driven by Tet-ON inducible *HRAS^G12V^* oncogene (28). Consistent with our sorting strategy, the expression patterns of *Cd80* and *Itga6* clearly distinguished three major clusters of cancer cells (Fig. 1E). In agreement with the bulk RNA-seq results, multiple immune modulatory genes such as *Cd276, Csf3, Il24* and *Il1a* (Fig. 1E) were selectively expressed in *Cd80*^+^*Itga6*^+^ cells, a population previously demonstrated to co-express the majority of stemness-associated genes (28).

### tSCs exhibit conserved immune modulatory programs in human SCCs

Next, to establish the human relevance of these findings, we analyzed single cell RNAseq data profiling cancer cell states in primary human CSCCs (41). As expected, the human SCC tumors display greater cellular heterogeneity, reflecting increased cellular plasticity. Nevertheless, based on the expression pattern of keratins and stem cell function genes (Fig. S2A), cancer cells could still be classified into tSCs, basal cells (comprising three major sub-clusters), and differentiated suprabasal cells (Fig. 1F). Notably, within the population defined as “tSCs”, which expresses the highest levels of *ITGA6* (Fig. 1F) and *KRT18* (Supplementary Fig. S2A), we detected clear stemness-associated signatures, including *SOX2* expression, together with a TGFβ responsive program marked by *VIM* (Supplementary Fig. S2A). Importantly, mirroring the observation in mouse SCCs, immune modulatory genes such as *CD80, CSF3, IL1A, IL24* and *PTGS2* are selectively expressed in this tSC population (Fig. 1F). A similar enriched expression pattern of these immune modulatory genes in stem cell-like cancer cells is also observed in human HNSCCs (Supplementary Fig. S2B), further supporting the concept that the immune modulation represents a shared stemness-associated molecular feature across SCC malignancies.

Taken together, by systematically comparing the expression patterns of known immune checkpoint ligands and secreted immune modulatory factors in distinct tumor populations in both CSCCs and HNSCCs, this study provides a comprehensive overview of the immune evasive and modulatory properties that are strongly enriched in stem cell-like tumor cells.

### Chromatin landscape shaping the tSC-specific immune modulatory program

Our comparative transcriptomic analysis demonstrated the selective upregulation of a defined cohort of immune modulatory genes while concomitantly downregulating immune stimulatory programs in tSCs. This coordinated pattern suggested the presence of an underlying regulatory mechanism that actively shapes immune selectivity in tSCs. We therefore hypothesized that stemness-associated transcriptional regulation shapes the specialized immune modulatory network in these cells. To test this hypothesis, we isolated CD80^+^ tSCs, CD80^-^ basal cells and differentiated tumor cells from DMBA/TPA induced SCCs and subjected these cells to Assay for Transposase-Accessible Chromatin using sequencing (ATAC-seq).

This analysis revealed distinct chromatin accessibility profiles across the three cancer cell populations (Fig. 2A and 2B). Importantly, chromatin accessibility closely mirrored the immune evasive states of tSCs, characterized by selective silencing at loci of key immune stimulatory genes such as *Cxcl9* and *Nlrp3*, together with increased accessibility at loci of immune suppressive or modulatory genes including *Cd276, Csf3, Tslp, Il1a,* and *Il24* (Fig. 2C and 2D). Motif analysis of differentially accessible chromatin regions between CD80^+^ tSCs and other tumor populations predicted the presence of binding elements for transcription factors activated by the TGFβ pathway (e.g., SMAD2/3), as well as regulators of inflammation (e.g., AP1 and STAT3) (Fig. 2E).

**Figure 2.**
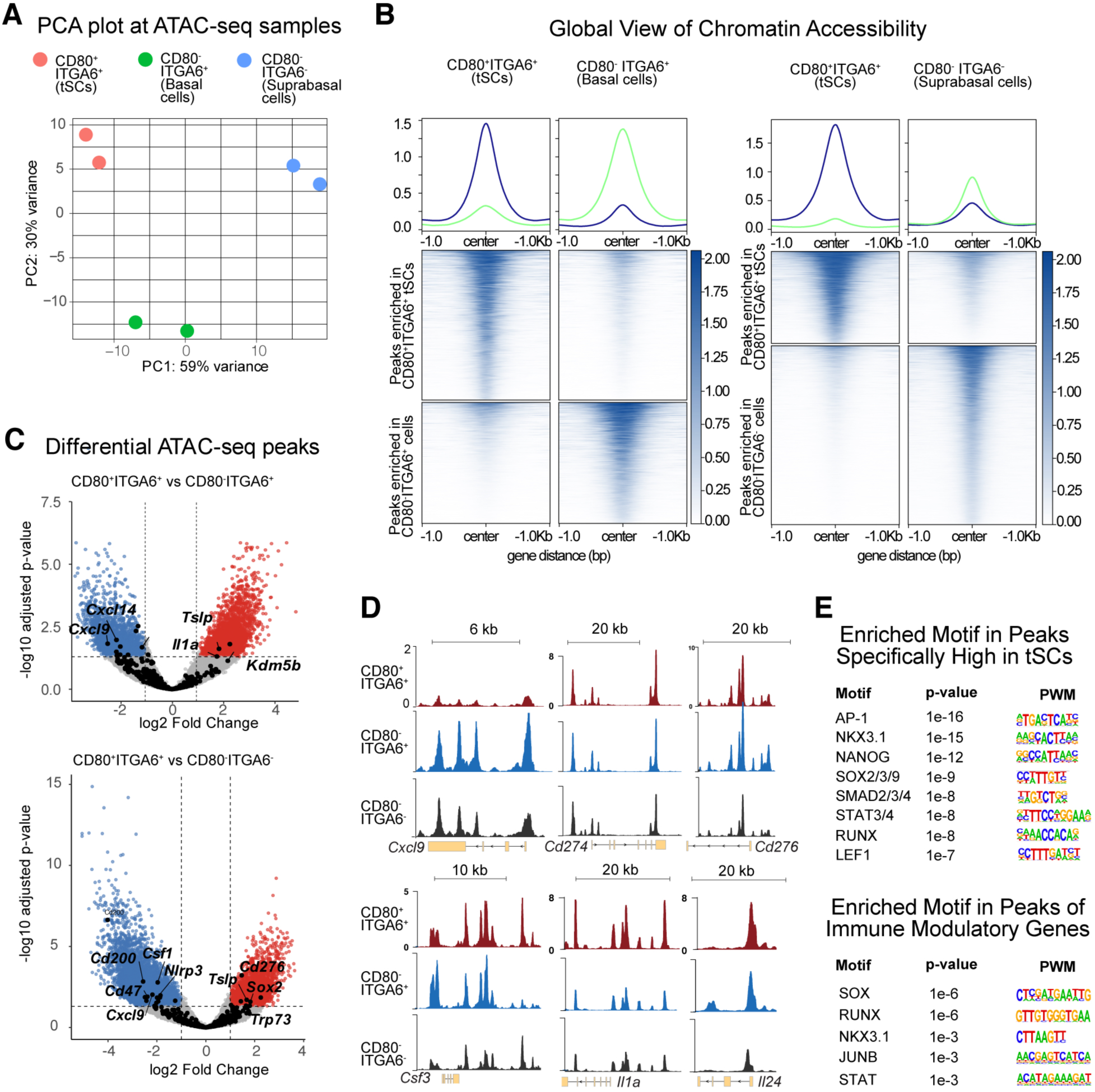
Distinct chromatin landscapes in tSCs are associated with their immune-modulatory programs. **A**, Principal component analysis (PCA) showing the distinct accessible chromatin profiles in tSCs compared to other tumor populations. **B**, Heatmaps and ATAC profile plots of differential peaks in CD80^+^ tSCs vs CD80^-^ basal cells or suprabasal cells. **C**, Volcano plots showing the differential peaks in CD80^+^ tSCs vs CD80^-^basal cells or suprabasal cells. **D**, IGV images showing the differential accessible chromatin regions near the representative immune modulatory genes in different tumor populations. **E**, Motif analysis on differential peaks between tSCs and other tumor populations (top) or the cohort of immune modulatory genes that are specifically enriched in tSCs (bottom).

Importantly, this analysis also revealed significant enrichment of SOX binding motifs within chromatin regions associated with immune-modulatory genes in tSCs (Fig. 2E). Among all the SOX family TFs, SOX2 emerged as a particularly compelling candidate, given its established roles in promoting tumorigenesis and maintaining tSCs identity in SCCs. Although SOX2 has not been classically implicated in immune regulation, recent work demonstrated that SOX2 is critical for SCC relapse following immunotherapy. This function is mediated, at least partially, through activation of FADS1, leading to altered lipid metabolism within the tumor microenvironment and suppression of interferon responses in TANs (26). Notably, the majority of immune modulatory genes identified in our analysis are preferentially enriched in cancer cells expressing high levels of SOX2 (Fig. 1F and Supplementary Fig. S2), suggesting that SOX2 may regulate broader immune modulatory activities in tSCs. We therefore next sought to investigate the functional significance of SOX2 in elevating immune-modulatory programs in tSCs.

### SOX2 broadly activates tSC-specific immune modulatory programs

To test the hypothesis that SOX2 can act as the master regulator of the immune modulatory program in tSCs, we first overexpressed *Sox2* in PDV cells, a highly immunogenic syngeneic mouse SCC cell line derived from the DMBA-treated keratinocytes isolated from C57/BL6 mice. These cells form stable SCC tumors only in immune deficient animals (42). When grafted on immune competent counterparts, PDV cells are rapidly cleared by the anti-tumor immune cells, suggesting that they have not evolved critical immune suppressive mechanisms. This feature makes PDV cells particularly suitable for evaluating the functional significance and underlying mechanisms of SOX2 in shaping the immune suppressive microenvironment in SCCs. One week post-grafting, prior to clearance of control tumors, SCC cells were isolated from vector control or *Sox2* overexpressing (OE) tumors and subjected to bulk RNA-seq. Transcriptome analysis demonstrated elevated expressions of multiple immune modulatory factors previously identified as enriched in tSCs, including *Csf3, Tslp, Il24, Il1a,* and *Mal,* specifically in *Sox2* OE PDV cells (Fig. 3A).

**Figure 3.**
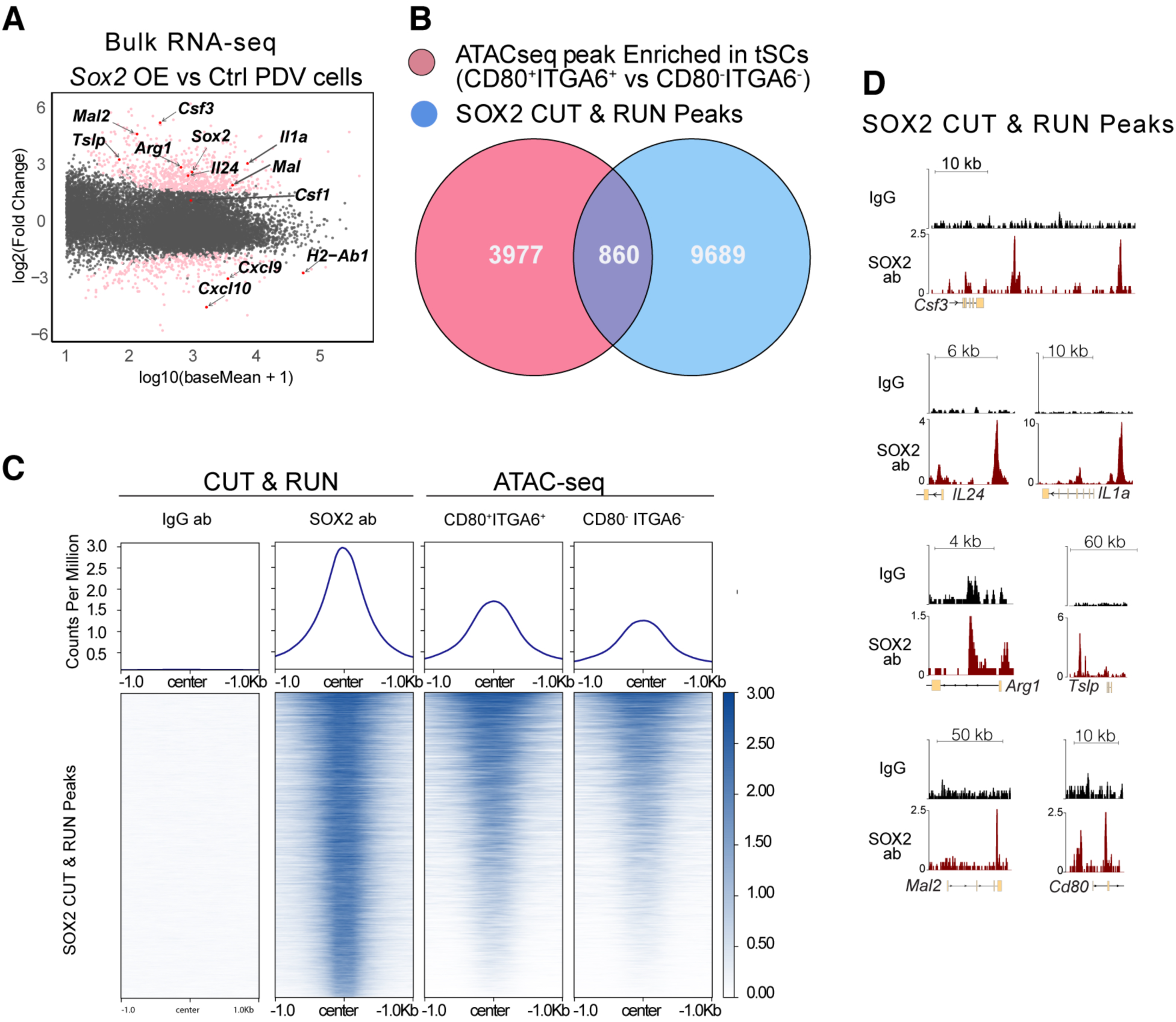
SOX2 directly activates immune modulatory programs in tSCs. **A**, MA plot showing the immune modulatory genes up-regulated following *Sox2* overexpression (<3 fold) in PDV cells. Genes with significant differential expression (adjusted p-value < 0.05) are shown as pink dots. **B**, Venn diagrams illustrating the overlap between chromatin regions selectively accessible in tSCs and regions bound by SOX2, as determined by SOX2 CUT&RUN (CNR). **C**, Heatmap of IgG control CNR, SOX2 CNR, and ATAC-seq signals in tSCs and differentiated cells across all SOX2-bound chromatin regions. **D**, IGV images showing SOX2 CNR occupancy at representative immune modulatory gene loci.

To confirm direct regulation of these immune modulatory genes by SOX2, we performed CUT & RUN sequencing using an antibody specific for SOX2. Comparison of SOX2-bound chromatin regions with those specifically accessible in tSCs revealed that at least 20% of the regions accessible in CD80^+^ tSCs are also directly bound by SOX2 (Fig. 3B). Consistent with this observation, tSCs possessed a greater number of accessible chromatin regions that are bound by SOX2 than other tumor populations (Fig. 3C). Importantly, SOX2 binding was detected at promoters and enhancers of the majority of tSC-specific immune modulatory genes (Fig. 3D). Taken together, these results establish SOX2 as a central transcriptional regulator that directly orchestrates the immune modulatory network in tSCs.

### Ectopic activation of SOX2 in normal epithelium is sufficient to expand and reprogram neutrophils

The identification of SOX2 as a regulator of the tSC-specific immune modulatory program prompted the development of complementary genetic models and functional assays to delineate how SOX2-orchestrated network enable epithelial cells to modulate immune populations. To isolate immunological effects attributable specifically to SOX2 activity, we ectopically induced *Sox2* expression in normal mouse epithelium in the absence of oncogenic transformation. For this purpose, we generated *Sox2* conditional overexpression (cOE: *K14CreER; R26-LSL-Sox2*) mice (Fig. 4A). As *Sox2* is transcriptionally silenced in the basal skin epithelium of adult mice, tamoxifen administration permits controlled induction of *Sox2* without activation of additional oncogenic pathways, thereby enabling direct assessment of its effects on immune cell modulation. To elicit immune responses that closely resemble those observed in cancer, we introduced cutaneous wounds in these mice. This experimental strategy is based on the well-established concept that cancers resemble “wounds that never heals” (43,44), reflecting shared biological processes between wound repair and tumor development, such as rapid epithelium proliferation and differentiation, extensive tissue remodeling and angiogenesis, and the recruitment of heterogenous immune cell populations that promote tissue regeneration (43,44). Thus, wound-induced inflammation provides a physiologically relevant system to evaluate the impact of ectopic SOX2 activation on immune cells regulation independently of tumor formation. Following tamoxifen administration, partial thickness wounds were generated in both control and *Sox2* cOE mice, and total CD45+ cells were isolated at the late stage of wound repair (day 7 post-wounding) for scRNA-seq and immune profiling (Fig. 4A). This time point was selected because the majority of wound recruited immune cells are short lived and are normally cleared by the late healing phase (i.e. Day 7), allowing epithelial stem cells to initiate tissue regeneration programs (43,45).

**Figure 4.**
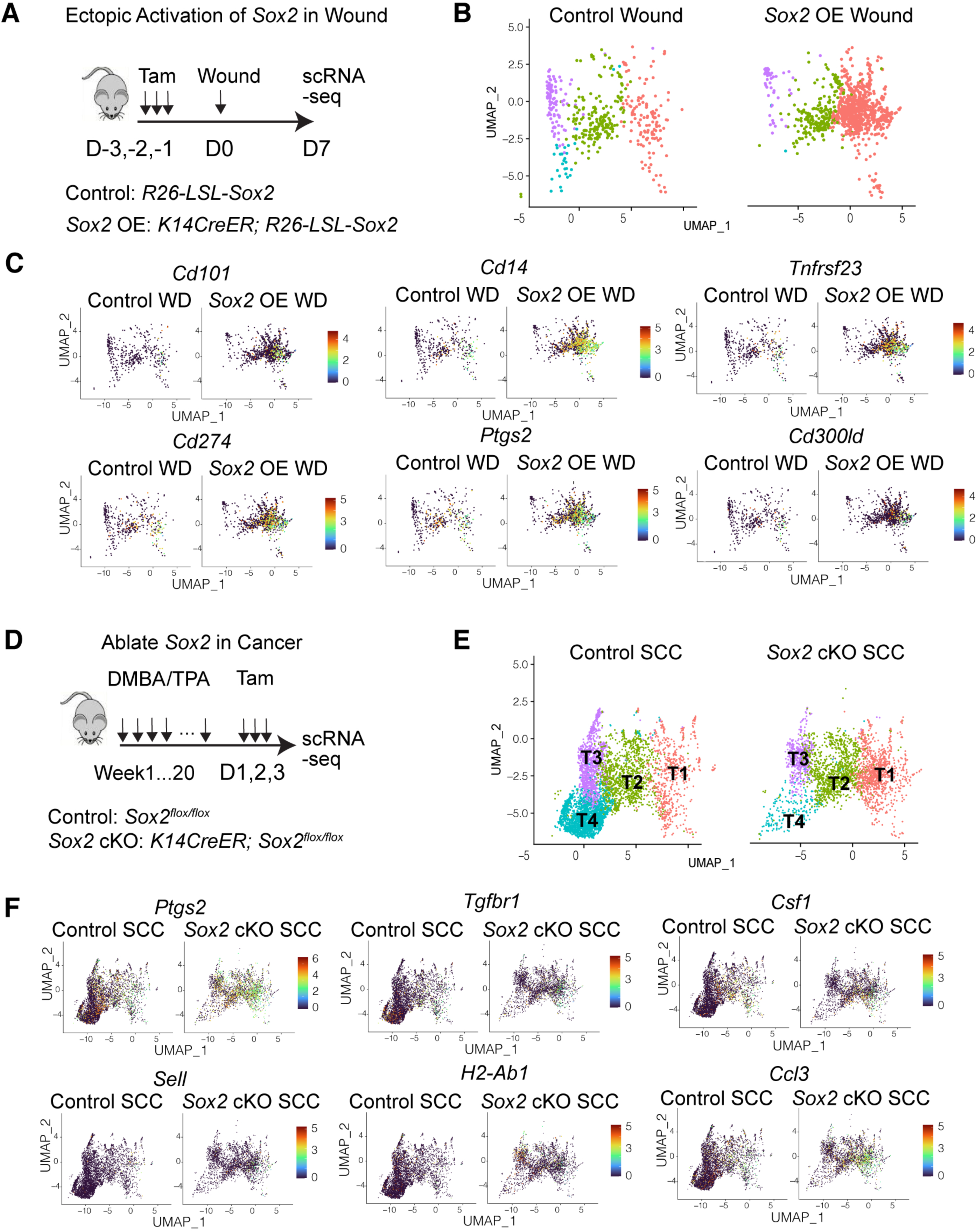
Genetic manipulation of SOX2 in skin epithelium remodels neutrophils in wound and cancer. **A**, Model schematics of ectopically activating *Sox2* in normal skin epithelium to assess how SOX2-induced programs influence the immune microenvironment during cutaneous wound repair. **B**, UMAP showing the increased accumulation of neutrophils in the wounded skin induced by ectopic activation of *Sox2*. **C**, UMAP showing the expression level of genes associated with neutrophil cell states and immune suppression in TANs with or without activating *Sox2* in untransformed epithelial stem cells during wound repair. **D**, Model schematics of *Sox2* ablation in DMBA/TPA-induced spontaneous SCCs to determine how loss of SOX2-driven programs in tSCs alters the immune microenvironment in the TME. **E**, UMAP showing the changes in the composition of neutrophil subpopulations following *Sox2* deletion in tSCs. **F**, UMAP showing differential expression of immune suppressive and immune-stimulatory genes expressed in TANs with or without deletion of *Sox2* in tSCs.

Analysis of the immune compartment revealed that wounds in *Sox2* cOE mice accumulated significantly higher numbers of neutrophils compared to control animals (Fig. 4B). Further transcriptional characterization of these neutrophils demonstrated induction of gene expression signatures characteristic of tumor associated neutrophils (TANs). In particular, elevated expression of CD14 (Fig. 4C), a defining marker of polymorphonuclear myeloid-derived suppressor cells (PMN-MDSCs)(46), was observed. PMN-MDSCs represent a specialized neutrophil state that mediates many of the immune suppressive and pro-tumor functions attributed to TANs (47,48). In addition, neutrophils accumulated in *Sox2* cOE wounds upregulate *Tfnrsf23* (gene encoding dcTRAIL) (Fig. 4C), a marker induced when neutrophils are deterministically programmed by the TME (49). Consistent with this phenotypic shift, neutrophils from *Sox2* cOE wound also upregulate multiple immune suppressive genes characteristic of TANs, such as *Cd274*, *Ptgs2* and *Cd300ld* (Fig. 4C). Together, these results indicate that SOX2 activates transcriptional programs that promote neutrophil persistence, thereby enabling acquisition of TANs-associated molecular features within an inflammatory environment.

### Ablating *Sox2* in tSCs decreases immune suppressive neutrophils in the TME

Next, we generated *Sox2* conditional knockout mice (cKO: *K14CreER; Sox2 ^flox/flox^*) (Figure 4D), enabling specific ablation of *Sox2* in the basal epithelium of the mouse skin. Spontaneous SCCs were then induced in both Cre negative control and *Sox2* cKO mice using DMBA/TPA. Following tamoxifen administration to induce Sox2 ablation, total CD45+ cells were isolated from SCC tumors and subjected to scRNA-seq to define alterations in the immune microenvironment resulting from blockade of SOX2-regulated programs in tSCs (Fig. 4D). Consistent with the role of SOX2-mediated programs in regulating neutrophil behavior observed in *Sox2* cOE wound model, scRNA-seq showed fewer neutrophils in *Sox2* cKO tumors (Fig. 4E).

Comparison of neutrophil cell states between control and *Sox2* cKO tumors identified pronounced alterations in TAN composition following *Sox2* ablation in tSCs. Consistent with previous reports (26), control SCC tumors harbored four major neutrophil subpopulations. Interestingly, *Sox2* deletion in tSCs specifically led to marked loss of both T3 and T4 neutrophils (Fig. 4E). Prior studies have shown that T3 and T4 TANs lack key markers of mature neutrophils, such as *Cd101* (49), while RNA velocity analyses demonstrated that T3 and T4 neutrophils differentiate towards the T1 and T2 neutrophil states (26). These findings suggest that TANs maintained by SOX2^high^ tSCs predominantly represent immature neutrophil population, which are known to possess potent immune suppressive activities in cancer contexts (47,48).

Importantly, comparative analysis of gene expression changes in TANs following *Sox2* ablation in tSCs revealed that neutrophil populations present in control tumors but depleted in *Sox2* cKO tumors expressed the highest levels of immune suppressive genes, such as *Ptgs2, Tgfbr1* and *Csf1* (Fig. 4F). In contrast, the residual TANs in *Sox2* cKO tumors exhibited increased expression of genes associated with anti-tumor neutrophil functions, such as Sell (50), MHCII genes (51), and T cell-recruiting chemokines (52). Collectively, these results define a previously unrecognized immune modulatory function of the SOX2-mediated tSC program that is essential for maintaining the immune suppressive neutrophils within the TME.

### SOX2 elevates CSF3 production from tSCs to expand neutrophils in the TME

The effects of genetic manipulation of *Sox2* on neutrophils are particularly intriguing, as these findings support the possibility that SOX2^high^ tSCs can shape their own immune suppressive niche. To further investigate this possibility, we performed immunofluorescence staining to assess the spatial distribution of neutrophils in DMBA/TPA induced spontaneous mouse SCCs. As expected, neutrophils were predominantly enriched at the tumor-stroma interface, coinciding with the location of SOX2^high^ tSCs (Fig. 5A).

**Figure 5.**
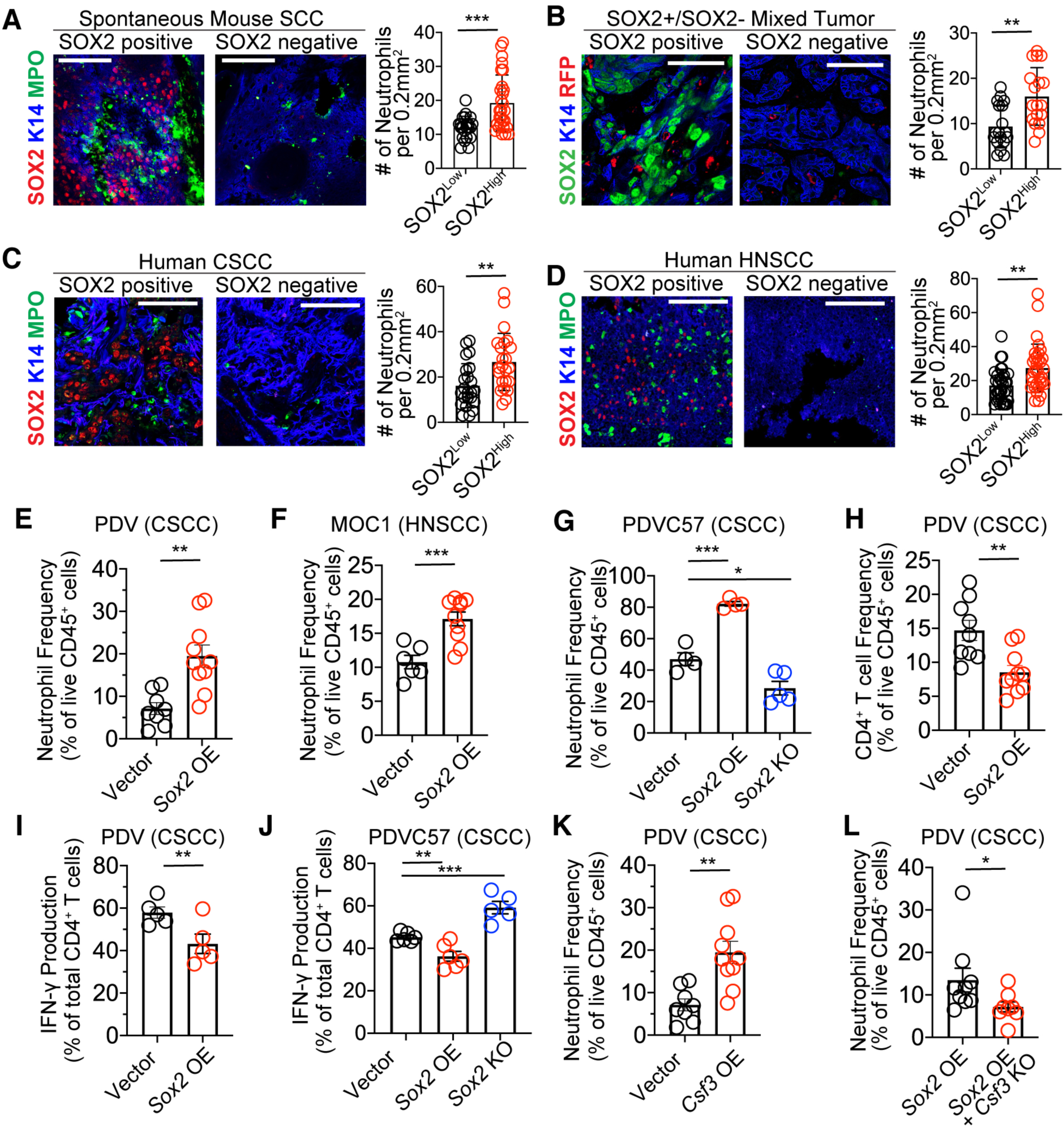
SOX2-driven CSF3 in cancer cells promotes neutrophil accumulation around SOX2^High^ tSCs. **A**, Representative IF images and quantification of the number of MPO^+^ TANs in the SOX2^High^ tSCs-concentrated (left panel) or the other SOX2^Low^ tumor regions (right panel) in mouse CSCCs. Scale bars: 100 μm. n = 25. **B**, Representative IF images and quantification of the number of RFP^+^ TANs per imaged area in SOX2^High^ (left) or SOX2^Low^ regions (right) when SOX2^Low^ and SOX2^High^ cells were mixed at 7:3 ratio for grafting on neutrophil-reporter mice. Scale bars: 50 μm. n = 18. **C** and **D**, Representative IF images and quantification of the number of MPO^+^ TANs per imaged area in the SOX2^High^ (left panel) or SOX2^Low^ tumor regions (right panel) in human CSCCs (**C**) or HNSCC (**D**). Scale bars: 100 μm. n = 30. **E–G**, Flow cytometry quantification of the neutrophils in CSCCs formed by immunogenic PDV cells (**E**), or HNSCCs formed by MOC1 cells (**F**), or immune resistant PDVC57 cells (G) with or without amplifying *Sox2*. n = 8 in (**E**), n=10 in (**F**), n=5 in (**G**). **H-J,** Flow cytometry quantification of CD4^+^ T cell infiltration (**H**), IFNγ production (**I** and **J**), in PDV (**H** and **I**) or PDVC57 cells (**J**) with or without amplifying or deleting *Sox2*. n = 10 in (**H**), n=5 in (**I**), n=6 in (**J**). **K** and **L**, Flow cytometry quantification of neutrophils in SCCs with or without amplifying *Csf3* (**K**) or silencing *Csf3* in *Sox2* overexpressing (OE) cells. n = 8 in (**K** and **L**). Bar graphs show representative results from one of the three repeats for each experiment and are presented as mean ± SEM. Student’s t tests (**A** to **D, E, F, H, I, K, L**) and one-way ANOVA followed by Tukey’s multiple-comparison tests (**G** and **J**) were used for statistical comparison. *p < 0.05; **p < 0.01, ***p < 0.001.

To directly test whether neutrophil distribution within the TME is shaped by SOX2-mediated programs, we designed a co-grafting assay. In this experiment, *Sox2* was amplified in PDVC57 cells, an immune resistant SCC line capable of forming tumors in immune competent mice. SOX2^High^ and SOX2^Low^ cells were then mixed at low ratio (3:7) and grafted onto neutrophil reporter ‘Catchup^IVM-red^’ mice (53). Analysis of neutrophil localization revealed that, despite representing a minority of the tumor cell population, SOX2^High^ regions exhibited significantly increased neutrophil accumulation (Fig. 5B). Importantly, a similar spatial enrichment of TANs in regions containing SOX2^High^ tSCs was also observed in primary human CSCCs (Fig. 5C) and HNSCCs (Fig. 5D), further confirming the clinical relevance of these findings.

To more quantitatively confirm the SOX2-mediated effects on neutrophil infiltration, we used flow cytometry to assess immune cells infiltration in SCC tumors with or without SOX2amplifiction. Consistent with our hypothesis, tumors with elevated SOX2 levels exhibited significantly increased infiltration of CD11b^+^Ly6G^+^ neutrophils in both CSCCs formed by PDV cells (Fig. 5E) and HNSCCs formed by MOC1 cells (Fig. 5F). To exclude the possibility that this difference is caused by conditions when parental PDV cells cannot establish stable tumors on immune competent mice,, we next mnipulated *Sox2* expression in PDVC57 cells, which form stable tumors even after Sox2 ablation. As expected, SOX2^High^ PDVC57 tumors again showed increased neutrophils infiltration, whereas *Sox2* KO tumors displayed a marked reduction in neutrophils (Fig. 5G). Importantly, our previous work demonstrated that, in the absence of immunotherapy, SOX2^High^ and SOX2^Low^ PDVC57 cells exhibit comparable tumor growth in immune competent mice (26). These results demonstrate that the observed differences in neutrophil infiltration are driven directly by SOX2-mediated immune modulation by cancer cells, rather than secondary effects of differential tumor growth. Increased neutrophil infiltration in SOX2^High^ tumors correlated with reduced CD4 T cell numbers (Fig. 5H and 5J, Supplementary Fig. S3A) and decreased production of anti-tumor cytokines, such as IFNγ (Fig. 5I and 5J, Supplementary Fig. S3A), while CD8 T cells were unaffected. Altogether, these data support the critical role of SOX2-driven tSC programs in promoting a neutrophil-mediated immune-suppressive TME.

Next, we explored the mechanisms by which SOX2^High^ tumor cells promote neutrophil expansion within the surrounding TME. Both transcriptome and CUT & RUN analysis identified *Csf3* as a key SOX2 regulated gene (Fig. 3) that is strongly enriched in tSCs (Fig. 1). The TME is known to recruit immature neutrophils, and elevated CSF3 levels have been shown to maintain TANs in an immature state (54). However, the cellular source of CSF3 within tumors remains elusive. To functionally test whether CSF3 is responsible for increasing the abundance of neutrophils in SOX2^High^ tumors, we either amplified *Csf3* in SOX2^Low^ SCC cells or deleted it in SOX2^High^ cells. As we expected, *Csf3* amplification in SOX2^Low^ cells recapitulated the neutrophil expansion seen with *Sox2* overexpression and resulted in increased neutrophil accumulation in the TME (Fig. 5K). Conversely, silencing *Csf3* in *Sox2* OE cells reduced neutrophil abundance (Fig. 5L).

Collectively, although cancer cells have been proposed to produce CSF3 to maintain immature neutrophil states after recruitment to the TME, our data demonstrate that enrichment of SOX2 in the tSCs further elevates CSF3 level that is critical for the expansion and maintenance of immature neutrophils within the TME. Through this mechanism, tSCs actively sculpt a myeloid cell-mediated protective niche that supports their survival and immune evasion.

### Loss of *Sox2* in tSCs alters the macrophage heterogeneity

In addition to the loss of specific neutrophil subsets following *Sox2* ablation, our single cell analysis also unveiled impaired maintenance of distinct tumor associated macrophage (TAM) populations. In spontaneous WT SCC tumors, five major macrophage clusters were identified (Fig. 6A). When *Sox2* was ablated in tSCs, macrophage clusters 3 (C3) and 4 (C4) macrophages were selectively depleted in the TME, while clusters 0 (C0) and 1 (C1) were expanded (Fig. 6A and 6B). Characterization of the depleted macrophage populations demonstrated that C4 macrophages express major markers of pro-tumor TAMs, including *Mrc1* (encoding CD206) and *Cd163*, as well as genes critical for TAM development such as *Maf*, and genes mediating immune suppressive functions, such as *Pf4* (Fig. 6C). In contrast, the persisting C0 and C1 macrophages exhibited increased expression of pro-inflammatory and anti-tumor associated genes, including class II MHC molecules (e.g. *H2-Ab1*), co-stimulation genes (e.g. *Cd80*), and inflammasome related genes (e.g. *Nlrp3)* (Fig. 6D). These findings suggest that the SOX2-mediated immune modulatory program in tSCs not only promotes expansion of the immature neutrophils but also maintains the pro-tumor macrophages within the TME.

**Figure 6.**
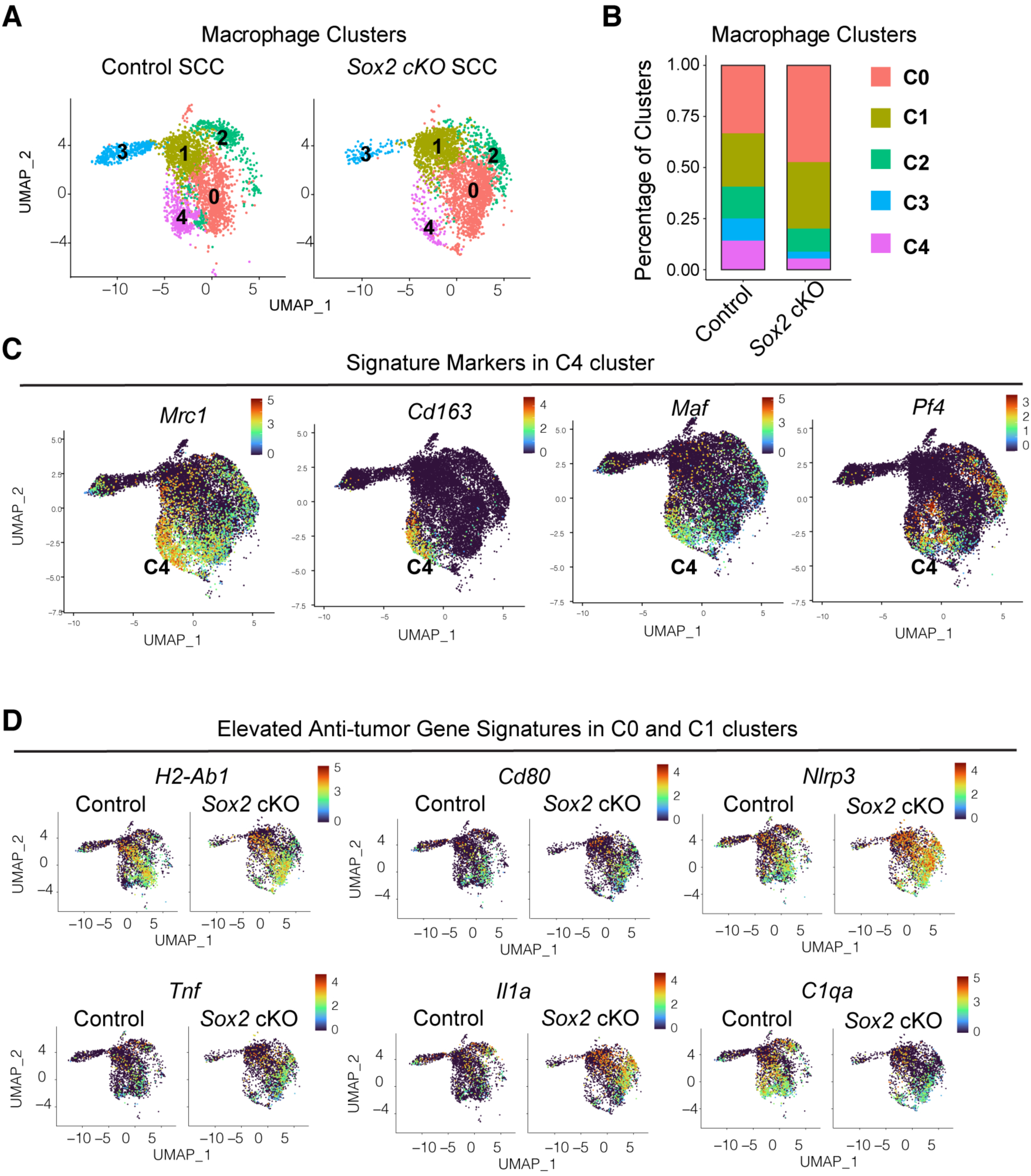
Silencing *Sox2* in tSCs selectively reduces CD163+ TAMs. **A** and **B**, UMAP (**A**) and stacked bar chart (**B**) showing the changes in the composition of macrophage subpopulations following *Sox2* deletion in tSCs. **C**, UMAP showing the expression of TAM signature genes enriched in the C4 macrophages subset which is reduced upon *Sox2* deletion in tSCs. **D**, UMAP showing expression level of various immune stimulatory genes in C0 and C1 macrophage populations that are expanded following *Sox2* deletion in tSCs.

### SOX2^high^ tSCs orchestrate the communication between TANs and TAMs

The dual role of SOX2^high^ tSCs in maintaining both TANs and TAMs are particularly notable. Given that SOX2 activates *Csf3* in tSCs, the mechanistic link between SOX2^high^ tSCs and immature neutrophils is clear. In contrast, transcriptomic profiling of tSCs and *Sox2* OE cancer cells, together with SOX2 CUT & RUN analysis, did not reveal an obvious candidate gene that could directly account for TAM regulation. It is well established that TAM development requires CSF1. Although previous studies proposed cancer cells as the major source of CSF1, yet SOX2 amplification did not significantly alter the expression of Csf1 in PDV cells. Interestingly, when we broadly searching for the source of CSF1 from our single cell analysis, we found that *S100a8^+^Csf3r^+^* neutrophils as the population expressing the highest levels of *Csf1* (Fig. 7A). This observation prompted us to explore whether immature neutrophils serve as an intermediary population that promotes TAM development and maintenance. To this end, flow cytometry quantification demonstrated increases in Ly6C^+^ macrophages in SOX2 OE SCC tumors (Fig. 7B and Supplementary Fig. S3B). Notably, silencing *Csf3* in SOX2^high^ SCC cells abolished the increase in macrophage accumulation (Fig. 7C). Given that scRNA-seq revealed a preferential loss of CD206+ TAMs in *Sox2* cKO tumors, we next specifically quantified CD206+ TAMs within the Ly6C+ macrophage population. Consistent with scRNA-seq findings, *Csf3* silencing in SOX2^high^ SCC cells selectively reduced the CD206+ TAMs subset (Fig. 7D). As *Csf3r* expression is largely restricted to neutrophils (Fig. 7A), these results indicate that the effects of *Csf3* ablation in SOX2^high^ cells are mediated through disruption of neutrophil-macrophage crosstalk rather than a direct effect on macrophages

**Figure 7.**
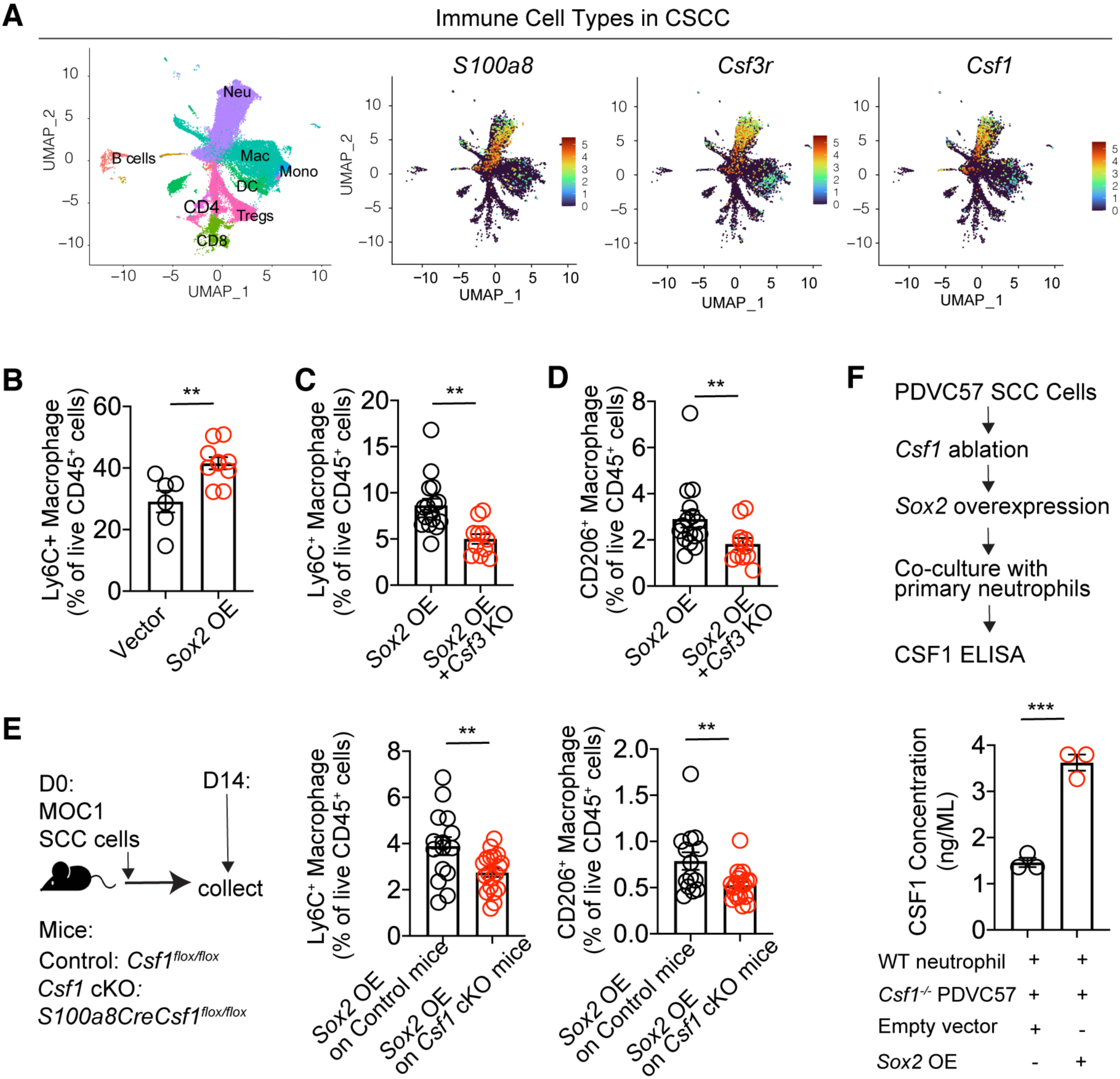
SOX2High tSCs promote the neutrophil-macrophage crosstalk to establish a protective niche. **A**. UMAP showing enriched expression of *Csf3r* and *Csf1* in *S1008a*^+^ TANs among all the immune populations in skin SCCs. **B-D**, Flow cytometry quantification of MHCII^+^CD64^+^Ly6C^+^ TAMs in SCCs with or without amplifying *Sox2* (**B**) or silencing *Csf3* in *Sox2* overexpressing (OE) cells (**C**) or CD206+ TAMs among MHCII^+^CD64^+^Ly6C^+^ macrophages in SCCs with or without silencing *Csf3* in *Sox2* overexpressing (OE) cells (**D**). n=6 or 9 in **B**, n=10 in **C**, n=10 in **D**. **E**. Experimental schemes, mouse models and flow cytometry quantification of Ly6C+ or CD206+ TAMs in SOX2^High^ SCCs grafted on mice with or without deleting *Csf1* specifically in neutrophils. n=15. **F**. Experimental schemes and ELISA quantification of CSF1 production from primary neutrophils when co-culture with in *Csf1*^-/-^*Sox2*^Low^ or *Csf1*^-/-^*Sox2*^High^ SCC cells. n=3. Bar graphs show representative results from one of the three repeats for each experiment and are presented as mean ± SEM. Student’s t tests (**C** to **F**) were used for statistical comparison. **p < 0.01.

To further assess this hypothesis, we generated the *Csf1* cKO mice (*S100a8Cre; Csf1 ^flox/flox^*) (Fig. 7E). SOX2^high^ SCC cells were then grafted onto either Cre negative control or the *Csf1* cKO animals. Strikingly, selective ablation of CSF1 production in neutrophils resulted in a significant reduction of CD206+ macrophages in SOX2^high^ tumors (Fig. 7E), providing support for a hierarchical regulatory axis where SOX2^high^ tSCs expand and program TANs, which in turn supply CSF1 to sustain pro-tumor TAM populations. To further strengthen this three-way crosstalk, we established an *in vitro* assay co-culturing either SOX2^Low^ or SOX2^High^ SCC cells with primary neutrophils (Fig. 7F). Since cancer cells can also produce CSF1, we employed CRISPR and generated *Csf1* KO SCC cells and then transduced these cells with either control vector or *Sox2*, enabling specific measurement of CSF1 originating from neutrophils (Fig. 7F). Interestingly, ELISA quantification showed that SOX2^high^ *Csf1^-/-^*SCC cells significantly elevated the CSF1 productions from neutrophils (Fig. 7F), further supporting our conclusion.

Taken together, this study identifies SOX2, a stem cell specific transcription factors, as a master regulator that shapes the immune modulatory activities of tSCs. Through this network, SOX2^high^ tSCs sustain elevated levels of immature neutrophils and CD206+ M2-like macrophages in their local environment, thereby sculpting a highly effective immune suppressive niche that confers superior protection from immune surveillance.

## DISCUSSION

Whether all cancers contain a subset of cells that function as tSCs remains under debate (55,56). However, even in malignancies where a clearly defined tSCs population cannot be identified, stemness is widely recognized as a core hallmark of cancer (57,58). Regardless of its cellular origin, acquisition of stemness endows cancer cells with the capacity to sustain the tumor growth, adapt to environmental stress, and survive diverse therapeutic interventions (59). Recently, enhanced immune resistance has emerged as an additional defining feature associated with cancer stemness (60,61). Despite this recognition, the mechanisms by which stem cell-like cancer cells acquire heightened immune resistance, and the reasons these cells preferentially survive immunotherapy while more differentiated tumor cells are eliminated, have not been systematically examined. Here, using SCCs which harbor a well-defined tSC population (28,31) as model systems, we integrated transcriptome profiling, chromatin landscape mapping, TF binding foot printing, genetic perturbation, and single-cell RNA sequencing, to obtain a global view of the immune evasive features and their underlying regulation in tSCs. This comprehensive analysis revealed that, compared to other cancer cells within the same tumors, a distinct population of CD80^+^ITGA6^High^ TGFβ responsive tSCs exhibits coordinated immune modulation characterized by reduced antigen presentation, diminished expression of proinflammatory chemokines, upregulated expression of immune suppressive ligands, and secretion of immune modulatory factors. Importantly, we identified the stemness-associated transcription factor SOX2 as a central regulator of this stem cell-specific immune evasive network.

The identification of SOX2 as a master regulator of immune modulatory network in tSCs is surprising. As a pioneer transcription factor that regulates embryonic stem cells, SOX2 is often silenced in adult tissue, but reactivated in many cancers upon acquisition of stemness (62,63). Although the pro-tumor functions of SOX2 are well established, prior studies have largely focused on its function in maintaining cancer cell self-renewal (62,63), leaving its broader roles in tumorigenesis insufficiently explored. Interestingly, SOX2 was found to be critical for dormant metastatic cancer cells to evade NK cell mediated killing (64). SOX2 also enables stem-like cancer cells to produce arachidonic acid to maintain the immune suppressive cell states of neutrophils (26). Although these processes can enhance the immune evasive capacity of cancer cells, these effects are still mediated by known SOX2 functions, such as modulating Wnt signaling (64) or regulating the lipid profile in the tumor (65,66). In contrast, the current study reveals that multiple immune modulatory factors enriched in tSCs (e.g. *Il24, Tslp,* and *Csf3*) are directly regulated by SOX2. Supporting its direct immune modulatory roles, we found that genetically manipulating the level of SOX2 remodels the immune microenvironment in both normal tissue and in the TME. We further dissected the functional consequences of disrupting the SOX2-CSF3 axis on the immune microenvironment. Additional SOX2-regulated immune modulatory genes identified in this study such as *Il24* or *Tslp*, remain to be functionally interrogated in future studies to define their individual and cooperative roles. Beyond SOX2, TGFβ-induced transcriptional regulation emerged as another potential contributor to the immune evasive features of tSCs. Although analysis of this pathway was beyond the scope of the current work, further investigation will be required to determine whether SOX2 and TGFβ signaling act synergistically to shape tSC-specific immune resistance (67). Given the limited systematic understanding of tSC-specific immune evasive programs, the comprehensive profiling presented here provides an essential framework for devising strategies to disrupt master regulatory mechanisms and downstream effectors of stemness-associated immune evasion.

Another intriguing finding of this study is the selective impact on specific subsets of myeloid cells following disruption of the SOX2-orchestrated tSC-specific immune modulatory network. Ablation of *Sox2* in tSCs of spontaneous mouse tumors resulted in the coordinated reduction of both immature TANs and CD206+ TAMs from the TME. These data uncover a previously unrecognized SOX2 driven immune modulatory program through which tSCs sustain immature neutrophil states in their local surroundings, which in turn support maintenance of CD206+ TAMs and establish stable myeloid crosstalk. The significant implication of this finding is consistent with recent insights onto how normal tissue stem cells adapt to inflammation (68). Under homeostatic conditions, tissue stem cells reside within immune privileged niches which protect them from inflammation-induced damages. However, how stem cells remain protected during injury repair remains incompletely understood, as tissue stem cells must exit their original niches, migrate into wound, and drive rapid tissue regeneration within a highly inflammatory environment. Our group and others have recently shown that immune modulation is an intrinsic feature of stem cells that plays an important role during wound repair. For example, hair follicle stem cells activate co-stimulatory molecules such as CD80 during wound healing to promote expansion of regulatory T cells, thereby facilitating termination of wound-induced neutrophil responses (45). This crosstalk allows stem cells to transiently sculpt a protective niche. Given the concept that ‘cancer is a wound that never heals’, analogous mechanisms are likely co-opted to shape the TME. The present study confirms this model and identifies a previously unknown three-way interaction among tSC, TANs, and TAMs. Importantly, tSCs emerge as the initiating population in this crosstalk and, through production of key cytokines, sustain the functional states of both TANs and TAMs.

Similar to normal tissue stem cells driving wound repair, tSC employ specialized transcriptional networks to construct a supportive niche within the TME. This niche enhances local immune suppression, enabling tSCs to evade immune surveillance while driving tumorigenesis. This extra protection provides a plausible explanation for the ability of tSCs to survive the enhanced anti-tumor immunity induced by immunotherapy, whereas more differentiated tumor cells in other regions of the TME are rapidly eliminated (28). Future work should be centered on testing strategies that disrupt crosstalk between tSCs and TANs, or between TANs and TAMs, as a therapeutic approach to abolish the immune privileged state of tSCs and prevent cancer relapse following treatment.

Finally, this study has important implications for understanding the biology of immune suppressive myeloid cells. Although the immune suppressive functions of immature TANs and CD206+ TAMs have been extensively characterized, their accumulation in the TME has largely been attributed to continuous chemokine-dependent recruitment, while the mechanisms governing their maintenance after entry into the TME remain poorly understood (69). CSF1 has been proposed as a key factor sustaining TAM populations within tumors (70), providing the rationale for therapeutic CSF1R blockade to reduce TAM abundance in the TME (71). Cancer cells have been considered to be the dominant source of CSF1 (72,73). Our findings challenge this paradigm and provide direct evidence that TANs are the primary producers of CSF1 that support TAM development in SCCs. Importantly, tSCs emerge as the upstream cell population that induces CSF1 production by TANs. To our surprise, this regulation is mediated through production of tSC-derived CSF3, raising several unresolved questions that warrant further investigation. For example, whether immature TANs are intrinsically wired to produce CSF1 and thereby support macrophage development once maintained within the TME. Thus, one possibility is that tSC-derived CSF3 primarily stabilizes TANs and maintains them in an immature state, thereby enabling sustained CSF1 production. Alternatively, CSF3 signaling may directly induce *Csf1* transcription in neutrophils. Distinguishing between these two equally plausible mechanisms was beyond the scope of the present study. Nevertheless, resolving these questions in future work will be important for advancing our understanding of myeloid cell development and for informing strategies aimed at normalizing myeloid cell functions as an alternative approach to controlling malignancy.

## METHODS

### Animal Experimental Models

*K14CreER* mice were generated by Dr. Elaine Fuchs lab and backcrossed to C57/BL6J background for more than ten generations (74). *Gt(ROSA)26Sor^tm1(CAG-Sox2,-EGFP)Blh^ (R26-LSL-Sox2)* mice were provided by Dr. Jianwen Que and have been described before (75). Wild-type C57BL/6J, *B6.Sox2^tm1.1Lan^/J* (*Sox2^flox/flox^*), B6.Cg-Tg(S100A8-cre,-EGFP)1Ilw/J (*S100A8-Cre*), C57BL/6-*Gt(ROSA)26Sor^tm1(HBEGF)Awai^*/J (*R26-LSL-DTR*) mice were obtained from The Jackson Laboratory. *Csf1*^flox/flox^ mice were provided by Dr. Jean X. Jiang. To induce *Sox2* conditional knockout, *K14CreER; Sox2^flox/flox^* mice were treated with daily intraperitoneal (i.p.) injection of 1 mg tamoxifen for three consecutive days. The same tamoxifen treatment regimen was used to induce *Sox2* expression in *K14CreER; R26-LSL-Sox2* mice. All mice were housed and bred under specific pathogen–free conditions in an Association for Assessment and Accreditation of Laboratory Animal Care (AAALAC)–accredited facility at The University of Chicago. All animal procedures were approved by the Institutional Animal Care and Use Committee (IACUC; protocol #72637) and conducted in accordance with the Guide for the Care and Use of Laboratory Animals.

### Cell Lines

The mouse skin squamous cell carcinoma (SCC) line PDV and PDVC57 and HEK293T cells were cultured in Dulbecco’s Modified Eagle Medium (DMEM) supplemented with 10% fetal bovine serum (FBS), 100 U/mL penicillin, 100 μg/mL streptomycin, and 2 mM L-glutamine. The mouse oral SCC line MOC1 was cultured in Iscove’s Modified Dulbecco’s Medium (IMDM)/F12 (2:1) supplemented with 5% FBS, 100 U/mL penicillin, 100 μg/mL streptomycin, 5 μg/mL insulin, 40 ng/mL hydrocortisone, and 5 ng/mL epidermal growth factor (EGF).

### Human samples

Fresh human CSCCs were collected immediately after surgical removal from patients at the University of Chicago Medicine under protocols approved by the Institutional Review Board (IRB) and in compliance with federal and state regulations and NIH guidelines. Tumors were then fixed, embedded in OCT, and cryo-sectioned for immunofluorescence imaging. Formalin-Fixed Paraffin-Embedded (FFPE) human HNSCC specimens were prepared and sectioned by the Human Tissue Resource Center at the University of Chicago and provided to us as slides.

### Tumor formation and Treatment

A two-stage chemical carcinogenesis protocol was used to induce skin SCCs in both female and male mice. At 8 weeks of age, the dorsal skin of various mouse strains was shaved and topically treated with 200 nM 7,12-dimethylbenz[a]anthracene (DMBA) once a week for 4 weeks to initiate tumorigenesis. Four weeks later, tumor promotion was induced by weekly (twice a week) applications of 200 µL of 35 µM 12-O-tetradecanoylphorbol-13-acetate (TPA) for 20 weeks. Oral squamous cell carcinoma (OSCC) was induced using 4-nitroquinoline 1-oxide (4NQO) administered in the drinking water as previously described¹. Briefly, mice received 4NQO (100 µg/mL) supplemented with 0.6% propylene glycol in the drinking water for 16 weeks, followed by regular drinking water for an additional 8–10 weeks to allow tumor development. Drinking water was replaced at least weekly and protected from light. Mice were monitored throughout the study for body weight and overall health. At experimental endpoints, tongues and oral tissues were harvested for downstream analyses and histopathological confirmation of SCC. For tumor transplantation, 2x10^6^ PDV mouse CSCC cells, 2x10^6^ MOC1 mouse HNSCC cells, or 5x10^5^ PDVC57 mouse CSCC cells were mixed with Cultrex Basement Membrane Extract, PathClear, Type 3 Matrigel (Bio-Techne) and injected subcutaneously. Tumors were allowed to grow for two weeks prior to immunotherapy. The tumor growth was measured twice a week, and tumor volumes were calculated using the formula: π/2 x length x width x height.

### Cell Sorting from Mouse Tumors

Tumors from DMBA + TPA-induced cutaneous SCCs, 4NQO-induced oral SCCs, and grafted mouse tumors were excised, minced, and enzymatically dissociated in RPMI-1640 (Gibco) containing 2 mg/mL collagenase type IV (Gibco) and 20 U/mL DNase I (Roche). Dissociation was performed for 30 min followed by 0.25% trypsin treatment for 10 min at 37°C for spontaneous SCCs, or for 60 min without trypsin for grafted tumors. Dissociated tissues were passed through a 70-µm cell strainer, subjected to red blood cell lysis using ACK buffer for 1 min, and resuspended in PBS. Single-cell suspensions were incubated for 15 min with biotin-conjugated antibodies against CD140a, CD45, CD11b, CD31, CD117, and CD64 (Biotin conjugated anti-CD11b (clone M1/70, Biolegend, Cat# 101204; RRID: AB_312787), Biotin conjugated anti-CD45 (clone 30-F11, Biolegend, Cat# 103104; RRID: AB_312969), Biotin conjugated anti-CD31 (clone MEC13.3, Biolegend, Cat# 102504; RRID: AB_312911), Biotin conjugated anti-CD117 (clone 2B8, Biolegend, Cat# 105804; RRID: AB_313213), Biotin conjugated anti-CD140a, rat monoclonal (clone APA5, Biolegend, Cat# 135910; RRID: AB_2043974), followed by incubation with 20 µL streptavidin-conjugated magnetic beads for an additional 15 min, to deplete stromal, hematopoietic, endothelial, and myeloid populations. Bead-bound cells were removed by magnetic separation to enrich for epithelial tumor cells. For isolation of ITGA^high^CD80⁺ tSCs and other tumor populations from spontaneous SCCs, enriched cells were stained with antibodies against CD49f and CD80 together with PE-conjugated streptavidin in FACS buffer. For grafted tumors, enriched cells were stained with PE-conjugated streptavidin alone. DAPI was added immediately prior to sorting to exclude dead cells. Live, lineage negative, ITGA6^high^CD80⁺, ITGA6^mid^CD80-, or ITGA6^-^CD80^-^ or GFP⁺ tumor cells were isolated using a BD Symphony S6 cell sorter and immediately processed for RNA extraction followed by RNA sequencing or ATAC-seq, as indicated.

### Partial-Thickness Wounding

Partial-thickness wounds were generated as previously described(76). Briefly, mice in the telogen phase of the hair cycle were shaved, and residual hair was removed using depilatory cream. Mice were anesthetized, and the dorsal skin was subjected to controlled abrasion using a Dremel drill head. Wound depth was standardized by applying a defined number of gentle passes, determined empirically by the appearance of uniform erythema with occasional pinpoint bleeding. This procedure selectively removes the epidermis and the upper portion of hair follicles, including most infundibulum and isthmus cells, while preserving the hair follicle bulge.

### Immune Cell Sorting from Tumors or Wounded Skin

Tumor samples were processed as described above. Wounded skin was processed using an adapted dissociation protocol(76). Briefly, excised tissue was incubated in cold PBS for 15 min to remove the scab, minced, and digested in RPMI-1640 (Gibco) containing Liberase (25 μg/ml; Roche) for 120 min at 37 °C. Digested tissue was passed through a 70-µm cell strainer, treated with ACK lysis buffer for 1 min to remove red blood cells, and resuspended in PBS. Single-cell suspensions were blocked with TruStain FcX (clone 93; BioLegend) in PBS supplemented with 5% normal rat serum and 5% normal mouse serum, stained with antibody cocktails at predetermined concentrations in FACS buffer (PBS containing 5% FBS, 5 mM EDTA, and 1% HEPES), and supplemented with DAPI to exclude dead cells. Live CD45+ cells were sorted from the single cell suspension of the tumor or wounded skin on a BD Symphony S6 cell sorter.

### Single Cell Suspension, Antibody Staining and Flow Cytometry Analysis

To profile immune cells from grafted tumors, tumors were excised, minced, and enzymatically dissociated in RPMI-1640 (Gibco) containing 2 mg/mL collagenase type IV (Gibco) and 20 U/mL DNase I (Roche) for 60 min at 37°C with gentle agitation. Dissociated tissues were filtered through a 70-µm cell strainer, subjected to red blood cell lysis using ACK buffer for 1 min, and resuspended in PBS. Cells were stained with Zombie Aqua Fixable Viability Dye (BioLegend) to exclude dead cells and then blocked with TrueStain FcX (clone 93; BioLegend) in PBS supplemented with 5% normal rat serum and 5% normal mouse serum. Single-cell suspensions were stained with a pre-optimized antibody cocktail diluted in FACS buffer (PBS containing 5% FBS, 5 mM EDTA, and 1% HEPES). The antibodies used include: APC/Cy7 anti-mouse CD45 (Clone 30-F11, Biolegend, Cat# 557659, RRID: AB_396774), APC anti-mouse CD206, (Clone C068C2, Biolegend, Cat# 141708, RRID: AB_10900231), AF488 anti-human/mouse TREM2 (Clone 237920, R&D Systems, Cat# FAB17291G, RRID: AB_3646994), FITC anti-mouse Ly6C (Clone HK1.4, Biolegend, Cat#128006, RRID: AB_1186135), BV605 anti-mouse TCR β Chain (Clone H57-597, Biolegend, Cat# 109241, RRID: AB_2629563), BV605 anti-mouse CD64 (Clone X54-5/7.1, Biolegend, Cat# 139323, RRID: AB_2629778), BV605 anti-mouse F4/80 (Clone BM8, Biolegend, Cat#123133, RRID: AB_2562305), PerCP/Cy5.5 anti-mouse Ly6C, rat monoclonal (Clone HK1.4, Biolegend, Cat#128011, RRID: AB_1659242), Alexa Fluor 700 anti-mouse CD45 (Clone 30-F11, Biolegend, Cat# 103128, RRID: AB_493715), PerCP/Cyanine5.5 anti-mouse/human CD11b (Clone M1/70, Biolegend, Cat# 101228, RRID: AB_893232), PE/Cyanine7 anti-mouse Ly-6G (Clone 1A8, Biolegend, Cat# 127618, RRID: AB_1877261), APC/Cy7 anti-mouse CD8b, rat monoclonal (clone YTS156.7.7, Biolegend, Cat# 126620, RRID: AB_2563951), PE/Cy7 anti-mouse CD4, rat monoclonal (clone GK1.5, Biolegend, Cat# 100422, RRID: AB_312707), BV711 anti-mouse TCR β Chain, armenian hamster monoclonal (clone H57-597, Biolegend, Cat# 109243, RRID: AB_2629564). PE-Granzyme B Monoclonal Antibody (clone GB11, Thermo Fisher, Cat# GRB04, RRID: AB_10853811), FITC anti-mouse IFN-γ, (clone XMG1.2, Biolegend, Cat# 505806, RRID: AB_315400). Flow cytometric analysis was performed on an LSRFortessa 4-15 HTS flow cytometer (BD Biosciences).

### Immunofluorescence staining and microscope imaging

Tumors freshly collected from mice or from human patients were fixed in 1% paraformaldehyde for 1 hour at 4°C and washed three times with cold PBS. After incubation in 30% sucrose at 4°C overnight, tumor tissues were embedded in OCT (Tissue Tek), frozen, and sectioned (10-15 μm). Cryosections were permeabilized, blocked, and stained with the following primary antibodies: Krt14 (chicken, 1:1000, BioLegend, Cat# 906004; RRID: AB_2616962), SOX2 (rabbit, 1:400, Cell Signaling Technology, Cat# #23064; RRID: AB_2714146), SOX2 (rat, 1:100, Thermo Fisher Scientific, Cat# 14-9811-82; RRID: AB_11219471), MPO (goat, 1:40, R&D, Cat# AF3667; RRID: AB_2250866),. The samples were then stained with corresponding secondary antibodies conjugated with Alexa Fluor 488, Rhodamine Red, or Alexa Fluor 647 (Jackson ImmunoResearch Laboratories) and imaged on Leica Stellaris 8 Laser Scanning Confocal microscope. For FFPE samples, slides were first deparaffinized by incubation at 60°C for 1 hour. Sequential washes (three times in xylene, followed by 100% ethanol, 95% ethanol, 75% ethanol, and distilled water (ddH2O), each for 10 minutes) were then used for rehydration. Antigen retrieval was performed using a low-pH antigen retrieval buffer (Thermo Fisher Scientific). Slides were then blocked at room temperature for 1 hour and incubated overnight at 4°C in a humidified chamber with the following antibodies: pan cytokeratin (rabbit, 1:100, Abcam, Cat# ab234297; RRID: AB_2895302), pan cytokeratin (mouse, 1:200, Novus Biologicals, Cat# NBP2-29429; RRID: AB_3068002), SOX2 (rat, 1:100, Thermo Fisher Scientific, Cat# 14-9811-82; RRID: AB_11219471), MPO (goat, 1:40, R&D Systems, Cat# AF3667; RRID: AB_2250866),. After primary antibody incubation, slides were washed three times with PBS, each for 5 minutes, and then incubated with secondary antibodies conjugated with Alexa Fluor 488, Rhodamine Red, or Alexa Fluor 647 (Jackson ImmunoResearch Laboratories). Slides were mounted using Prolong Gold Antifade Mountant (Thermo Fisher Scientific), carefully avoiding air bubbles during coverslip placement. Fluorescent images were acquired using a Leica Stellaris 8 Laser Scanning Confocal microscope and quantified on ImageJ.

### Neutrophil Isolation and Culture

Neutrophil isolation was performed using protocols adapted from published methods(77). For isolation from grafted tumors, tumors were excised, minced, and enzymatically dissociated in RPMI-1640 (Gibco) containing 2 mg/mL collagenase type IV (Gibco) and 20 U/mL DNase I (Roche) for 60 min at 37°C with gentle agitation. Dissociated tissues were filtered through a 70-µm cell strainer, and red blood cells were lysed using ACK buffer for 1 min. Single-cell suspensions were resuspended in 8 mL of 40% Percoll and overlaid onto 3 mL of 70% Percoll in 15-mL conical tubes. Samples were centrifuged at 900 × g for 30 min at room temperature without brake. Cells at the interface were collected, washed twice, and resuspended in FACS buffer. Cells were blocked with TruStain FcX (clone 93; BioLegend) in PBS supplemented with 5% normal rat serum and 5% normal mouse serum for 10 min, followed by incubation with 20 µL biotin-conjugated anti-Ly6G antibody for 15 min and 20 µL streptavidin-conjugated magnetic beads for an additional 15 min. After four washes with FACS buffer, neutrophils were resuspended in RPMI-1640 for downstream experiment. For isolation from bone marrow, femurs were harvested, epiphyses removed, and marrow flushed with RPMI-1640 using a syringe, followed by filtration through a 40-µm cell strainer. For spleen-derived neutrophils, spleens were mechanically dissociated and filtered through a 40-µm strainer. Red blood cells from bone marrow and spleen samples were lysed using ACK buffer for 1 min, after which neutrophils were isolated using the same magnetic enrichment procedure described above. Freshly isolated neutrophils were then cultured in RPMI-1640 media with 10% FBS, 100 U/mL penicillin, 100 μg/mL streptomycin, 55 μM beta-mercaptoethanol (GIBCO), 50 U/mL GM-CSF (Biolegend).

### CRISPR-Mediated Gene Knockout

Gene knockouts were generated using the lentiCRISPRv2 system. Single-guide RNAs (sgRNAs) targeting Csf3 (CACCGACTTAAGCAGGAAGCTTCG) were designed using CRISPick (Broad Institute) and cloned into the lentiCRISPRv2 vector (Addgene #52961). Lentiviral particles were produced by transfecting HEK293T cells with the lentiCRISPRv2 construct together with the packaging plasmids psPAX2 and pMD2.G. Viral supernatants were collected 48 h post-transfection, filtered through a 0.45-µm filter, and used to transduce target cells in the presence of 10 µg/mL polybrene. Transduced cells were selected with puromycin (2 µg/mL) for 3 days to enrich for successfully transduced populations. Single-cell clones were generated by limiting dilution. Genomic DNA was extracted using the Quick-DNA Microprep Kit (Zymo Research), and the sgRNA target regions were PCR-amplified and sequenced to confirm gene editing.

### Bulk RNA Sequencing and Analysis

Total RNA from FACS-sorted cells was isolated using the Quick-RNA Microprep Kit (Zymo Research) according to the manufacturer’s instructions. RNA-seq libraries were prepared using the NEBNext Single Cell/Low Input RNA Library Prep Kit for Illumina (New England Biolabs) and sequenced on an Illumina NovaSeq platform. Raw FASTQ files were adapter-trimmed and quality-filtered using Cutadapt (v3.2). Transcript abundances were quantified against the mouse mm10 reference genome (GENCODE vM24) using Kallisto (v0.44.0). Gene-level counts were summarized, and differential expression analysis was performed using DESeq2 (v1.30.0) in R (v4.1.1). Genes with Benjamini–Hochberg–adjusted p values < 0.05 were considered significantly differentially expressed.

### ATAC-seq Library Preparation and Data Analysis

Tumor cells were isolated by FACSs as described above. 100,000 cells were used for ATAC-seq library preparation. ATAC-seq libraries were generated using the Active Motif ATAC-seq Assay according to the manufacturer’s instructions. Briefly, sorted cells were pelleted, washed with cold PBS, and lysed to isolate nuclei. Nuclei were incubated with pre-indexed assembled Tn5 transposomes in tagmentation buffer at 37°C for 30 min to simultaneously fragment chromatin and insert sequencing adapters. Following tagmentation, DNA was purified and PCR-amplified to generate sequencing-ready libraries. Libraries were size-selected, quantified, and assessed for quality prior to sequencing. ATAC-seq libraries were sequenced on an Element Biosciences Aviti platform using paired-end reads. For data analysis, raw sequencing reads were adapter-trimmed and quality-filtered using Cutadapt (v3.2)(78). Clean reads were aligned to the mouse reference genome (mm10) using Bowtie2 (v2.3.4.3), and PCR duplicates were removed with Picard (v2.18.7). For visualization, BAM files from biological replicates were merged, and ATAC-seq signal tracks were generated using deepTools(79) and visualized with the Gviz R package(80). Peaks were called using MACS2 (v2.2.7.1)(41), and peaks from all samples were combined and merged using BEDTools (v2.30.0)(81). Read counts for each merged genomic region were quantified using featureCounts (v1.5.3)(82). Differential chromatin accessibility analysis was performed using DESeq2, with significance determined based on Benjamini–Hochberg–adjusted p values.

### CUT&RUN Data Processing and Visualization

CUT&RUN sequencing data were obtained from GEO (accession GSE278426). Raw FASTQ files were adapter-trimmed and quality-filtered using Cutadapt (v3.2)(78) and aligned to the mouse reference genome (mm10) with Bowtie2 (v2.3.4.3). PCR duplicates were removed using Picard (v2.18.7), and peaks were identified using MACS2 (v2.2.7.1)(41). For visualization, BAM files from biological replicates were merged, and normalized signal tracks were generated using bamCoverage (deepTools v3.5.6)(79) with counts per million mapped reads (CPM) normalization. BigWig files were visualized using the Gviz R package.

### Single-Cell RNA-Seq Data Analysis

The FASTQ files were first processed by spipe.v1.1.1 provided by Parse Biosciences, using mm10 as a reference. Counts data were imported to analyzed using Seurat(83). Low-quality cells were filtered based on gene number and mitochondrial content. We then used the standard Seurat pipeline by running NormalizeData, FindVariableFeatures, and ScaleData. For the first round of clustering of total immune cells, principal component analysis (PCA) was performed and the top 50 PCs with a resolution = 0.6 were applied. RunUMAP was used to visualize the data. Cell types were then annotated using SingleR (V1.7.1) (84) package with ImmGen data as reference. Neutrophils or macrophages were subset, and standard pipeline was applied to neutrophil or macrophage population by running FindVariableFeatures, and ScaleData. For clustering the neutrophils or macrophages, Principal component analysis (PCA) was re-performed and the top 40 PCs with a resolution = 0.2 were applied. For visualization, RunUMAP was used. For gene signature analysis, we performed pesudobulk analysis by randomly grouping cells into 3 pseudo replicates. Gene counts were summarized and DESeq2 R package (v1.30.0)(85) in R (v4.1.1) was used for differential gene analysis.

### Quantification and Statistical Analysis

Data are presented as mean ± SEM or mean ± 95% confidence interval, as indicated. Statistical significance was determined using two-tailed Student’s *t* test, Mann–Whitney *U* test, or one-way or two-way ANOVA, as specified in the figure legends. Statistical analyses were performed using GraphPad Prism 9. Experiments were conducted in an open-label manner. Significant difference between two groups were noted by asterisks (* p < 0.05; ** p < 0.01: *** p < 0.001).

## Data Availability

New single cell RNA sequencing, bulk RNA sequencing, ATAC-sequencing, and CUT & RUN-seq data reported in this paper have been deposited at Gene Expression Omnibus (GEO) and are publicly available as of the date of publication. Accession numbers include (GSE278435, GSE278426). Previously published single cell RNA-seq data were also analyzed and the accession numbers include GSE144239. Tumor growth, microscopy data and flow cytometry data reported in this paper will be shared by the lead contact upon request. This paper does not report original code and any additional information or data in this paper will be available from the lead contact upon request

## Author Contributions

Y.M, W.G. and D.L. conceptualized the study, designed the experiments, interpreted the data, and wrote the manuscript. W.G. and D.L. performed most experiments and analyzed the data with the assistance of X.H., J.L., and B.N. B.D. collected human CSCC samples, M.L., A.P., E.I., A.R. N.A. collected and provided the HNSCC samples.

## Acknowledgments

We thank J. Stanisavic, S. Fisher in the Miao lab for assistance; C. Ciszewski at the Human Disease & Immune Discovery Core Facility at the UChicago for conducting FACS sorting; H. Shah (Metabolomics Platform at the UChicago Comprehensive Cancer Center) for measuring AA in TIF; P. Faber (Genomics Core at the UChicago) for sequencing and raw data processing; Animal Resources Center at UChicago for assisting animal work. This study was supported by Y.M.’s Start-up fund from UChicago, Cancer Research Foundation Breakthrough Board, Cancer Center Support Grant number (P30 CA014599), Pilot grants from The University of Chicago Medicine Comprehensive Cancer Center, grants to Y. M. from NIH (R00CA237859, R01CA285786), American Cancer Society, V Foundation, American Association for Cancer Research, and The Cancer Research Foundation.

