## Supplemental information for "Integrated analysis of stemness-associated immune modulatory circuits in squamous cell carcinomas"

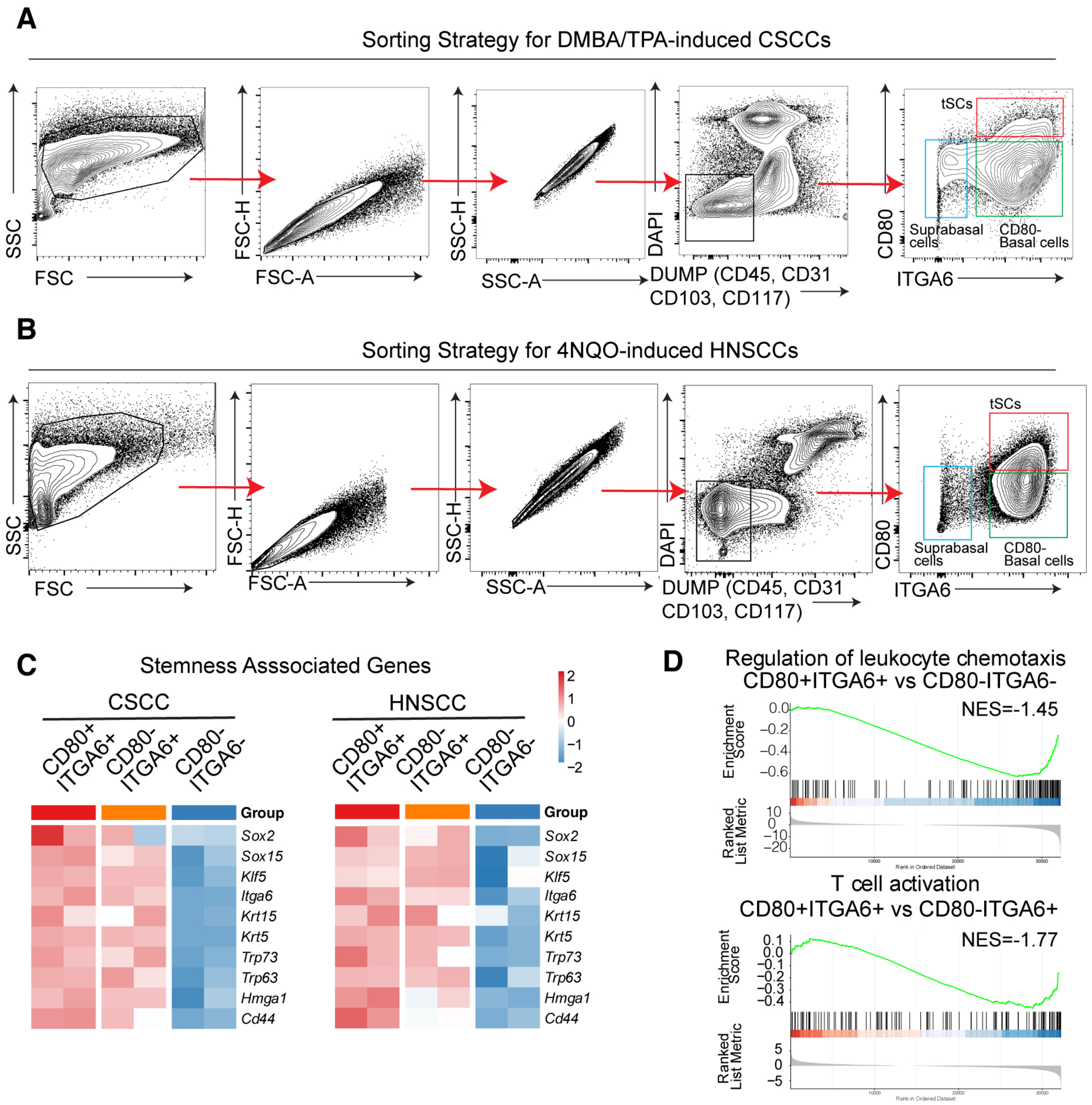


**Supplementary Figure S1 Transcriptome analysis of tSCs in SCCs**

**A** and **B**, Sorting strategies to isolate different tumor populations from mouse CSCCs (**A**) or HNSCCs (**B**). **C**, Heatmap showing the differentially expressed stemness-related genes in CD80$^+$ITGA6$^+$ tSCs, CD80$^-$ITGA6$^+$ basal cells, and CD80$^-$ITGA6$^-$ suprabasal cells in CSCCs and HNSCCs. Scale: relative expression. **D**, Leading edge plots showing the downregulated immune stimulatory pathways in CD80$^+$ITGA6$^+$ tSCs, compared to CD80$^-$ basal cells or ITGA6$^-$ suprabasal cells in CSCCs

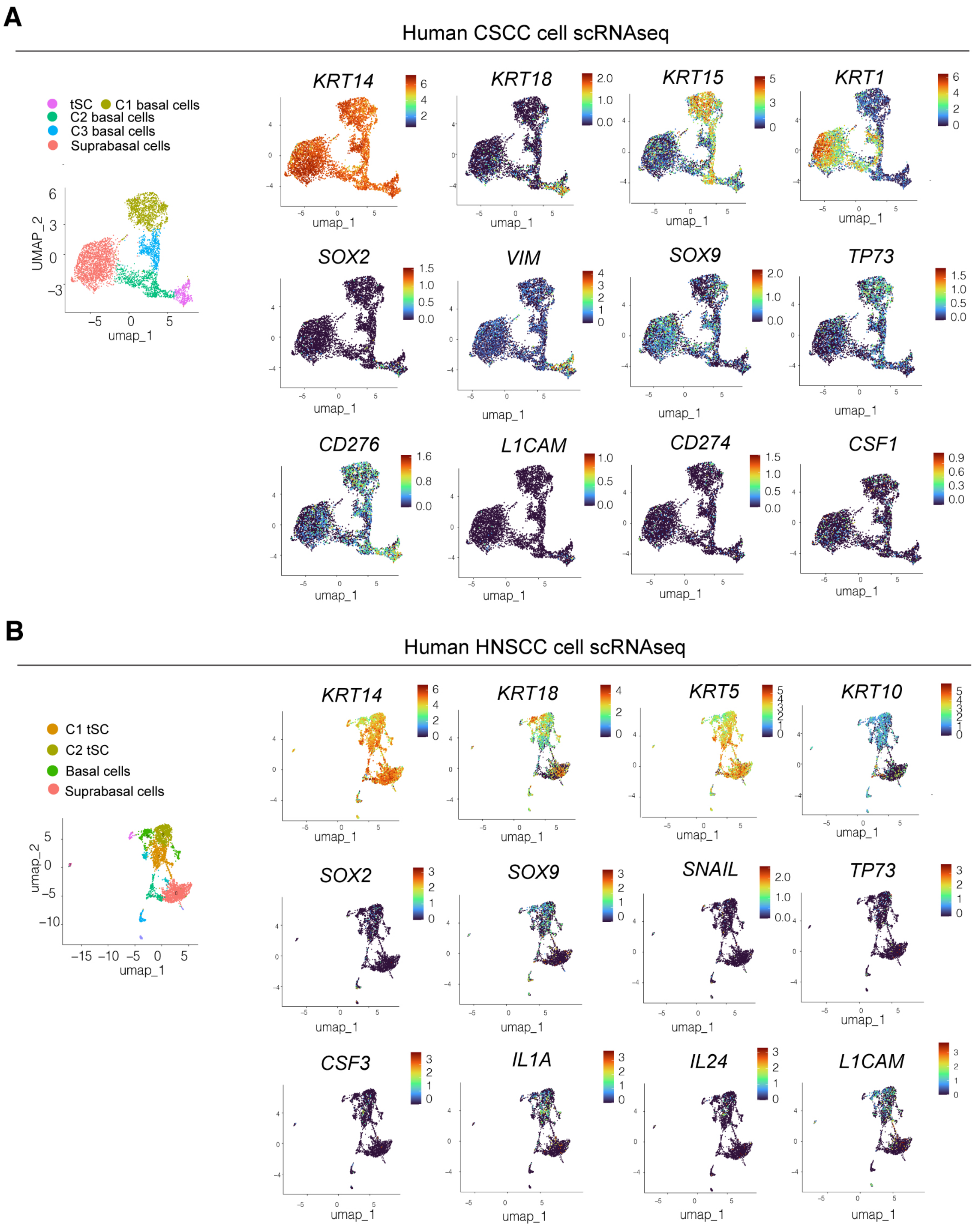


**Supplementary Figure S2. Single cell analysis of human SCC signatures.**
**A** and **B**, UMAP showing the signature and immune modulatory genes expressed in various cancer cell clusters in human CSCCs (**A**) or human HNSCCs (**B**). Scale: relative expression.

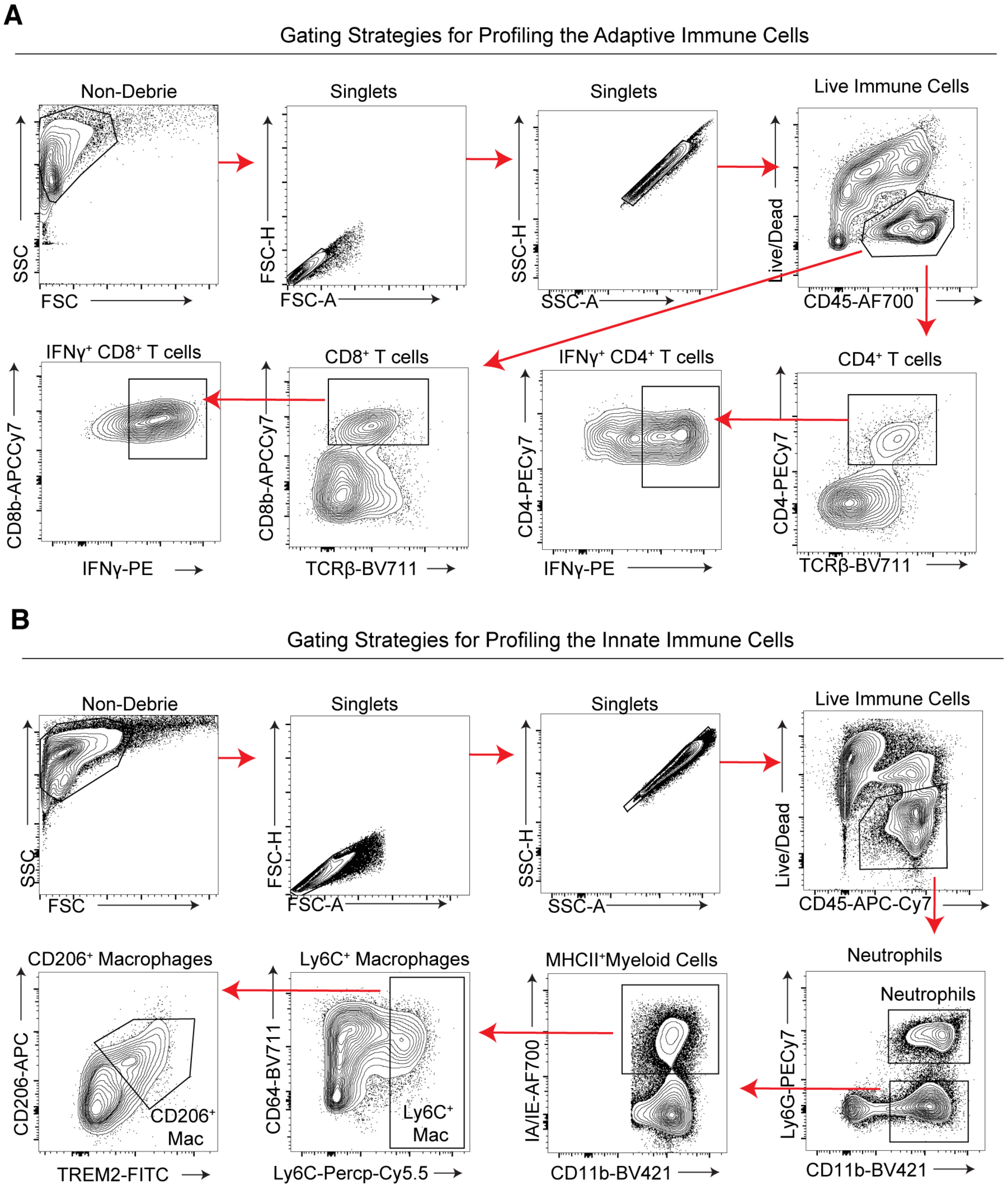


**Supplementary Figure S3 Immune profiling in grafted mouse SCCs**

**A** and **B**, Gating strategies to profile the adaptive immune cells and anti-tumor cytokine production (**A**), or innate immune cells (**B**) in grafted SCCs.
